# Spine-neck electrical bottlenecks tune temporal precision and inhibitory gating in cortical pyramidal neurons: A connectomics-based biophysical study

**DOI:** 10.64898/2026.02.16.706119

**Authors:** Netanel Ofer, Sapir Shapira, Idan Segev

## Abstract

Dendritic spines are nano-scale compartments that host the majority of excitatory synapses on cortical pyramidal neurons (PNs); hundreds of spines/PN also receive an inhibitory synapse (dually-innervated spines, DiSs). Using analytic theory and detailed biophysical models of ∼2,000 densely reconstructed spines, we show that recurrent spine-neck narrowing substantially increases spine-neck resistance (*R*_*neck*_), and that elevated *R*_*neck*_ accelerates spine-head voltage dynamics, shortening spinous postsynaptic potentials by up to ∼3-fold. *R*_*neck*_ improves the tracking of high-frequency synaptic inputs and strongly modulates *Ca*^2+^ signaling and the potency and temporal precision of inhibitory gating in DiSs. This work identifies *R*_*neck*_ as a dynamic “knob”, directly linking spine ultrastructure to information processing and plasticity in cortical PNs and circuits, yielding testable experimental predictions. Our “biophysics of connectomics” paradigm naturally raises computational-oriented questions, including how inhibitory “gates” in dendritic spines expand context-dependent computations, implement single-cell and network-level routing, and enable selective encoding of precise temporal patterns.

## Introduction

Dendritic spines densely cover the dendritic tree of cortical pyramidal neurons (PNs). They have fascinated neuroscientists since the pioneering work of Ramón y Cajal, who meticulously drew these tiny, highly variable protrusions, consisting of a bulbous head connected via a thin, cable-like neck emerging from the parent dendrite (Cajal, 1911). Cajal hypothesized that spines held essential secrets about neuronal function (DeFelipe and Jones, 1992). The advent of electron microscopy (EM) in the mid-20th century subsequently revealed the detailed ultrastructural organization of dendritic spines at nanometer resolution (Gray, 1959; Peters and Kaiserman-Abramof, 1970). Quantitative studies showed that PNs typically host several thousand dendritic spines (Elston et al., 2001; Benavides-Piccione et al., 2024) and that the vast majority of excitatory synapses in PNs target spines (Larkman, 1991). Notably, a significant fraction (up to 15%) of PN spines also receive a second, inhibitory synapse – the dually innervated spines (DiSs) (Kubota et al., 2007; Chen et al., 2012; Chiu et al., 2013; Müllner et al., 2015; Bloss et al., 2016; Tremblay et al., 2016; Boivin and Nedivi, 2018; Gemin et al., 2021; Tullis and Bayer, 2024).

Subsequent experimental and theoretical work established that the spine head acts as a tiny electrical and biochemical compartment that can be enriched with voltage-gated Na^+^ and *Ca*^2+^ conductances, as well as AMPA and NMDA receptors. Owing to their small dimensions and partial electrical isolation from the parent dendrite via the thin neck, spines may exhibit high input impedance that amplifies local excitatory inputs, promotes cooperativity among spiny inputs, and supports local *Ca*^2+^-dependent plasticity processes (Rall, 1974; Segev and Rall, 1988; Araya et al., 2007, 2014; Doron et al., 2017; Harris and Kater, 1994; Nimchinsky et al., 2002; Schiller et al., 2000; Yuste and Denk, 1995; Grunditz et al., 2008).

Of particular functional importance is the spine neck, whose nanometer-scale diameter imposes a large axial resistance (*R*_*neck*_). Theory and sophisticated experimental measurements indicate that, depending on neck dimensions, *R*_*neck*_ can reach the *G*Ω range (Rall, 1974; Svoboda et al., 1996; Grunditz et al., 2008; Araya et al., 2014; Harnett et al., 2012). Rall (1974) proposed that the neck length/diameter may change in an activity-dependent manner, thus acting as a dynamic plastic “device” controlling both the magnitude of excitatory synaptic potential (EPSP) at the spine-head as well as the transfer of synaptic current to the parent dendrite and soma (see also (Crick, 1982)). Consistent with this view, extensive experimental work has shown that dendritic spines are structurally plastic across the cortex and hippocampus; changes in spine geometry and connectivity were observed over minutes to months; these dynamics covary with learning and memory processes (Engert and Bonhoeffer, 1999; Yuste and Bonhoeffer, 2001; Trachtenberg et al., 2002; Knott et al., 2002; Holtmaat and Svoboda, 2009; Yuste, 2010, 2013; Takeuchi et al., 2014; DeFelipe, 2015; Berry and Nedivi, 2017; Eyal et al., 2018).

Only recently, through the connectomics efforts, have large datasets of thousands of EM-reconstructed spines become available with sufficient resolution to bridge the ultrastructure of neurons and circuits to their computational implications. Serial EM reconstructions, in hippocampus and cortex, complemented by modern *in vivo* imaging, reveal substantial spine-to-spine and branch-to-branch variability in head and neck geometry and spine-density (Harris and Stevens, 1989; Kasthuri et al., 2015; Ofer et al., 2021, 2022; Benavides-Piccione et al., 2025). This rich data enables an EM-based, biophysical framework, which we hereby call *“biophysics of connectomics”*, to ask how nano-scale neck geometry governs the temporal precision of excitatory/inhibitory integration within individual spines. We hereby show that repeated, often extreme, neck narrowing creates electrical bottlenecks that results with large *R*_*neck*_. In addition to affecting the magnitude of spinous inputs, we found that *R*_*neck*_ strongly modulates spine-head voltage dynamics, shaping the operating time-window of EPSPs and IPSPs interaction, and strengthening the inhibitory gating in DiSs. Together, these results identify nanoscale bottlenecks in spine necks as tunable, geometry-defined control parameters that set both the time-window for synaptic integration and the strength of local inhibitory veto at single dendritic spines. We discuss the functional implications of these results for context-dependent computations, information routing, and the selective encoding of precise temporal patterns.

## Results

### Large variability in spine neck morphology quantified from serial EM reconstruction

This study is based on the analysis of a large-scale EM dataset comprising 3D reconstructions of 2,074 dendritic spines from the apical tree of layer 6 pyramidal neurons from the mouse somatosensory cortex (Kasthuri et al., 2015). This analysis provided the most detailed measurements of the highly irregular spine neck geometry, thus offering new estimates for the spine neck resistance (*R*_*neck*_). Consequently, using EM-based compartmental modeling, we assessed the impact of *R*_*neck*_ on local synaptic signaling, focusing on its impact in the temporal domain.

Figure 1A shows an exemplar dendritic branch consisting of 132 dendritic spines, with a density of 2.62 spines/*µm*. Two representative spines (orange and blue) are highlighted for further analysis. The EM-based high-resolution triangular mesh reveals the tortuous shape and pronounced variability in neck diameter for both spines, particularly for the orange (longer) spine (Figure 1B). The spine head is depicted in green and the central axis of the spine neck is marked by the black dots. The neck of the orange and blue spines was reconstructed using 202 and 142 sections, respectively (see **Methods**). Two EM slices of the orange spine are also shown; the top right demonstrates an extreme restriction at the narrowest point of the spine neck, with a diameter of approximately 30*nm*.

**Figure 1.**
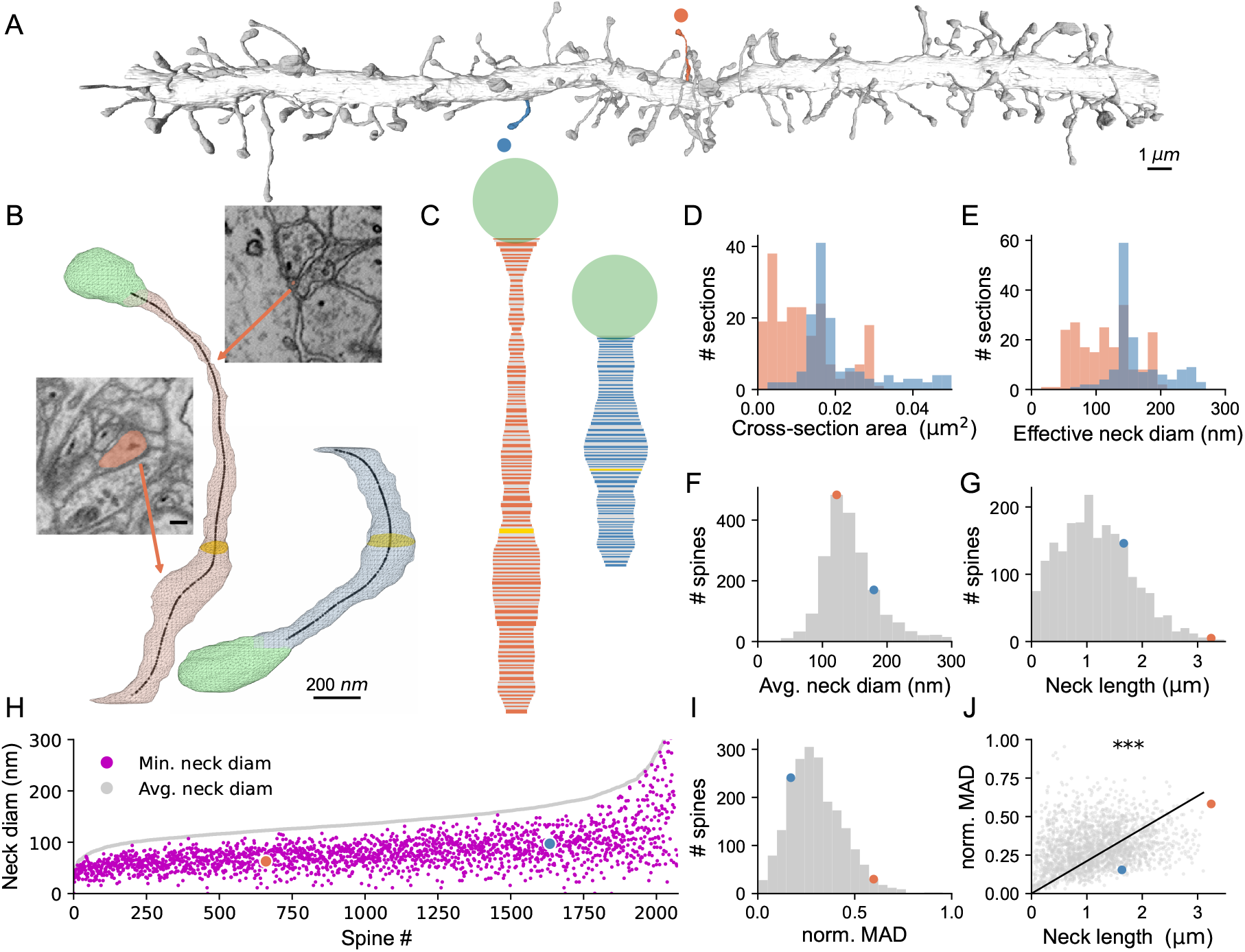
Highly heterogeneous spine neck geometry with exceedingly constricted neck diameter revealed by serial EM reconstruction. **A.** Representative EM-reconstruction of apical dendritic branch of L6 pyramidal neuron from mouse somatosensory cortex. Two exemplar dendritic spines (orange and blue) are highlighted. **B**. Triangle-mesh reconstructions of the two exemplar spines; spine heads are shown in green, and spine necks in orange and blue, respectively. Black dots trace the central line (skeleton) of the spine neck; representative cross-section ellipses are marked in yellow. Two EM images taken along the “orange” spine neck (arrows) illustrate thin (top right, diameter ∼30*nm*) and thick regions (lower left, average diameter ∼270*nm*). Scale bar for both EM images is 100*nm*. **C**. Effective diameter, *d*, profiles along the spine necks for the two spines in B (n = 202 and 142 sections for orange and blue spines, respectively, see **Methods**). Yellow sections correspond to the yellow sections in B. **D, E**. Distributions of cross-sectional areas and effective diameters, respectively, along the necks of the two spines shown in B. **F**. Distribution of the average spine neck diameter across n = 2,074 spines, calculated as the weighted mean of *d* per-section. The two exemplar spines are shown by orange and blue circles. **G**. Distribution of spine neck lengths across all reconstructed spines. **H**. Scatter plot comparing average (gray) and minimum (magenta) spine neck diameters across all reconstructed spines. Spines are sorted by their average neck diameter. **I**. Distribution of the normalized median absolute deviation (MAD) index (see **Methods**). **J**. Correlation between spine neck length and MAD index (R = 0.85, p < 0.001, Pearson correlation test). Data from (Kasthuri et al., 2015).

To visualize diameter variability along the spine neck, we plotted the effective diameter, *d*, of each section along the spine neck (Figure 1C); *d* was calculated from the cross-sectional area, *A*, per section, 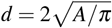. Figures 1D and 1E depict the distributions of *A* and *d*, respectively, for the two exemplar spines. Neck geometry differs substantially between the two spines: the orange spine exhibits highly variable neck diameters, whereas the blue spine shows a more uniform, cylindrical-like profile. To characterize the spine-neck morphology over all our dataset, we computed, for each spine, the weighted average diameter (*d*_*w*.*avg*_) using Equations 1 and 2,

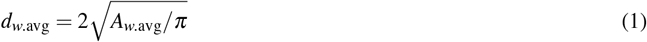

where,

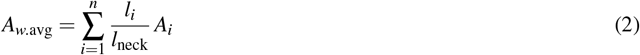

*l*_*i*_ and *A*_*i*_ are the length and cross-sectional area of the *i*-th section along the spine neck, respectively, and *l*_*neck*_ is the total length of the neck. Across the full dataset, the average neck diameter was 148 ± 55 nm (mean ± SD); median, 137 nm; central 99% range, [55 - 409 nm] (Figure 1F). Average spine neck length was 1.19 ± 0.69 *µm* (mean ± SD); median, 1.11 *µm*; central 99% range, [0.05-3.38 *µm*] (Figure 1G; exemplar spines shown in orange and blue).

To highlight variability in spine-neck diameter along individual spines, we compared the average (gray) and minimum (magenta) neck diameters for each spine across all reconstructed spines (Figure 1H). Most spines exhibit considerable diameter narrowing along their neck; some spines show extremely thin segments along their neck (on the order of tens of nanometers). To quantify the intra-spine irregularity, we computed the normalized median absolute deviation (MAD) index (Figure 1I; see **Methods**). The majority of spines displayed pronounced variability in neck diameter (normalized MAD: 0.30 ± 0.14, mean ± SD; median 0.28), with multiple constrictions and widenings along the spine neck. Figure 1J demonstrates that diameter irregularity and neck length were positively correlated (R = 0.85, p < 0.001, Pearson correlation test), indicating that longer spines tend to exhibit a greater variability in neck diameter.

In conclusion, from serial EM reconstruction of 2,074 spines, we found that neck diameters were highly non-uniform, with frequent constrictions below 100*nm*. The normalized MAD index shows that only a minority of spine necks have a near-cylindrical shape. These fine-scale variations necessitate EM-based estimation of spine geometry, setting the stage for modeling how the ultrastructure of the spine neck affects a key parameter – the spine neck resistance (*R*_*neck*_). In turn, we will explore the effect of *R*_*neck*_ on the temporal dynamics of synaptic signaling at the spine head.

### New EM-based estimates of spine neck resistance (*R*_*neck*_)

Our estimates of *R*_*neck*_ from anatomical measurements are based on the cable equation (Rall, 1974),

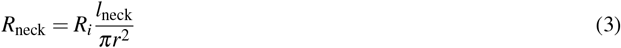

where *R*_*i*_ is the specific axial resistivity (in Ω*cm*) of a cylindrical cable, *l*_*neck*_ and *r* are the spine’s neck length and radius, respectively. Note that neck resistance depends linearly on the neck’s length and reciprocally on its diameter squared.

To estimate *R*_*neck*_ from EM data, we applied two methods. In the “average” method, we used the weighted-average cross-sectional area, *A*_*w*.*avg*_, along the spine neck,

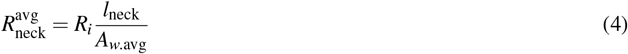

In the “section-by-section” method, *R*_*neck*_ was computed for each section individually, and the contributions of all *n* sections composing the spine neck were then summed.

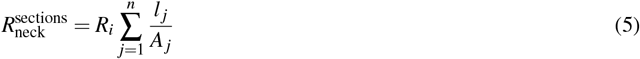

where *l* _*j*_ and *A*_*j*_ are the length and cross-sectional area of the *j*-th section. Unlike the “average” method (Equation 4), the “section-by-section” method captures local geometrical variations and therefore provides a more accurate estimate of *R*_*neck*_.

To illustrate the differences between the two methods, we used a simplified model of a spine neck composed of two cylindrical segments with radii *a < b* and lengths *l*_*a*_ and *l*_*b*_, respectively (Figure 2A). The total neck length was held constant, while *l*_*b*_*/l*_*a*_ and *b/a* varied. Both methods yield identical values in the case of a cylinder (black dashed diagonal), but they diverge when the two cylindrical segments differ in diameter. Figure 2A shows that 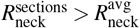 with discrepancy increasing as the diameter ratio, *b/a*, increases (*b/a* = 2 vs. *b/a* = 5, dark and light blue, respectively). This is proven analytically in **Appendix A**. for the general case, where the spine neck is composed of several cylindrical segments of different diameters.

**Figure 2.**
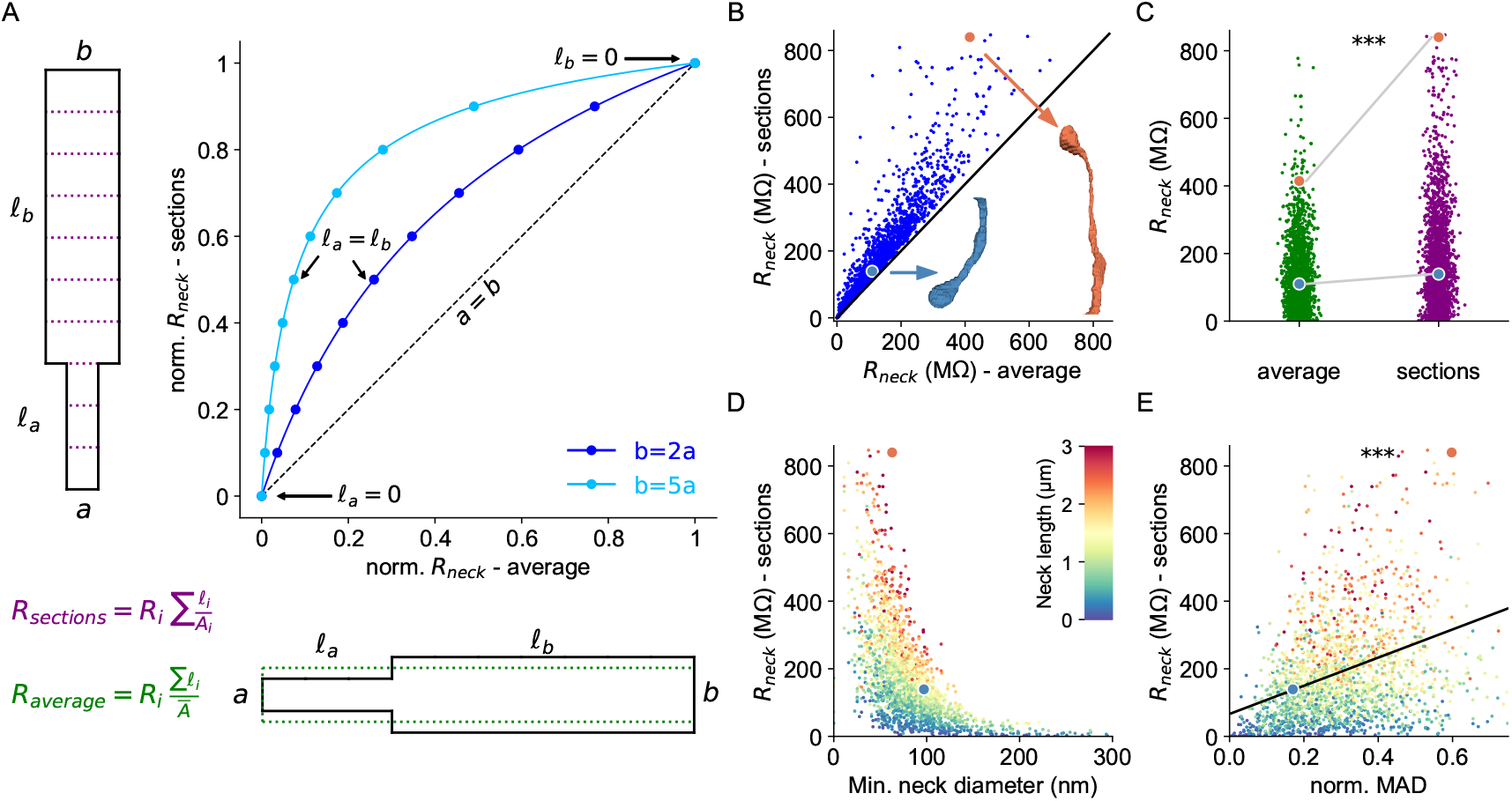
Irregular spine-neck diameter elevates spine-neck resistance. **A.** Comparison of two approaches for computing spine-neck resistance (*R*_*neck*_): the weighted average-diameter method (*R*_*average*_; x-axis, lower equation in green) versus the section-by-section method (*R*_*sections*_; y-axis, upper equation in purple). The toy geometry comprises two cylindrical segments with diameters *a* and *b*, lengths *l*_*a*_ and *l*_*b*_, and fixed total length *l* = *l*_*a*_ + *l*_*b*_. The dotted green cylinder illustrates the single cylinder with weighted average diameter used for computing *R*_*average*_; this diameter was used to compute the expected *R*_*neck*_ in the x-axis. The dotted purple lines indicate the discrete sections used for computing *R*_*sections*_ (y-axis). Dark- and light-blue curves correspond to *b/a* = 2 and *b/a* = 5, respectively; points along each curve depict various *l*_*a*_*/l*_*b*_ ratios. Dashed black diagonal indicates the case where *a* = *b*. **B**. *R*_*sections*_ versus *R*_*average*_ for n = 2,074 EM-reconstructed spines. Orange and blue points (also shown in C-E) correspond to the two exemplar spines in Figure 1A-C. **C**. Distribution of *R*_*neck*_ across all reconstructed spines using the average (green) versus section-by-section (purple) methods. The former method significantly underestimates *R*_*neck*_ (paired t-test, p < 0.001). **D**. *R*_*sections*_ as a function of minimum neck diameter. Color encodes neck length (scale inset); longer necks tend to exhibit smaller minimum diameters. **E**. Positive correlation between MAD index for neck-diameter irregularity and *R*_*sections*_ (R = 0.32, p < 0.001, Pearson correlation test), indicating that more irregular necks have higher resistance. *R*_*i*_ = 150Ω*cm* for all *R*_*neck*_ computations.

Applying both methods to estimate *R*_*neck*_ over the full dataset of 2,074 spines confirmed that the “average” method systematically underestimates *R*_*neck*_ (Figure 2B). The two exemplar spines are indicated by the orange and blue circles (inset). Figure 2C shows that the section-by-section approach (purple) yields, on average, 50% larger *R*_*neck*_ values (191.6 ± 182 *M*Ω, mean ± SD; median 150.3 *M*Ω; central 99% range, 2 - 918 *M*Ω) compared to the “average” method (green; 132.8 ± 105.2 *M*Ω, mean ± SD; median 109 *M*Ω; central 99% range, 2 - 569 *M*Ω; paired t-test, p < 0.001). With *R*_*i*_ = 150 Ω*cm*, some spines reached *R*_*neck*_ values approaching 1 *G*Ω (Figure 2C, see **Discussion**).

To better understand how the irregular geometry of the spine neck affects *R*_*neck*_, we plotted the correlation between the minimal neck diameter and *R*_*neck*_. Figure 2D shows that *R*_*neck*_ strongly depends on the minimum neck diameter; *R*_*neck*_ tends to increase sharply in spines with pronounced neck narrowing, implying that thin sections along the neck act as ‘bottlenecks’ that contribute disproportionately to the overall neck resistance. Noteworthy is that most cases of small minimal diameter are associated with large neck resistance, which typically occurs in longer spine necks (color-coded in Figure 2D). Furthermore, *R*_*neck*_ was significantly and positively correlated with the MAD index, confirming that greater spine irregularity is associated with larger neck resistance (Figure 2E, R = 0.32, p < 0.001, Pearson correlation test).

Together, the results of Figure 2 demonstrate that conventional estimates of *R*_*neck*_ based on average neck diameter substantially underestimate the true *R*_*neck*_ value. In contrast, the “section-by-section” method, grounded in automated morphometric analysis of EM-based 3D spine reconstructions, provides more accurate *R*_*neck*_ estimates. Below, we show that *R*_*neck*_ strongly impacts the temporal dynamics of spinous EPSPs.

### Spine neck resistance shapes temporal dynamics at the spine head

In the reconstructed ∼20*µm* dendritic branch shown in Figure 3A, spines exhibit striking heterogeneity in neck dimensions, leading to an estimated *R*_*neck*_ ranging from ∼100*M*Ω (blue spines) to ∼1*G*Ω (red spines). This broad range of *R*_*neck*_ values implies that nearby spinous synapses can experience markedly different voltage responses and different levels of filtering of synaptic inputs. To study the effect of *R*_*neck*_ on the dynamics of spinous EPSP, we simulated a single spine receiving excitatory synaptic input emerging from a long passive cylindrical dendritic cable (schematic in Figure 6B; see **Methods**). An AMPA-like excitatory transient conductance was delivered to the spine head (Figure 3B, dotted line); the resulting EPSPs were recorded at the spine head for different *R*_*neck*_ values.

**Figure 3.**
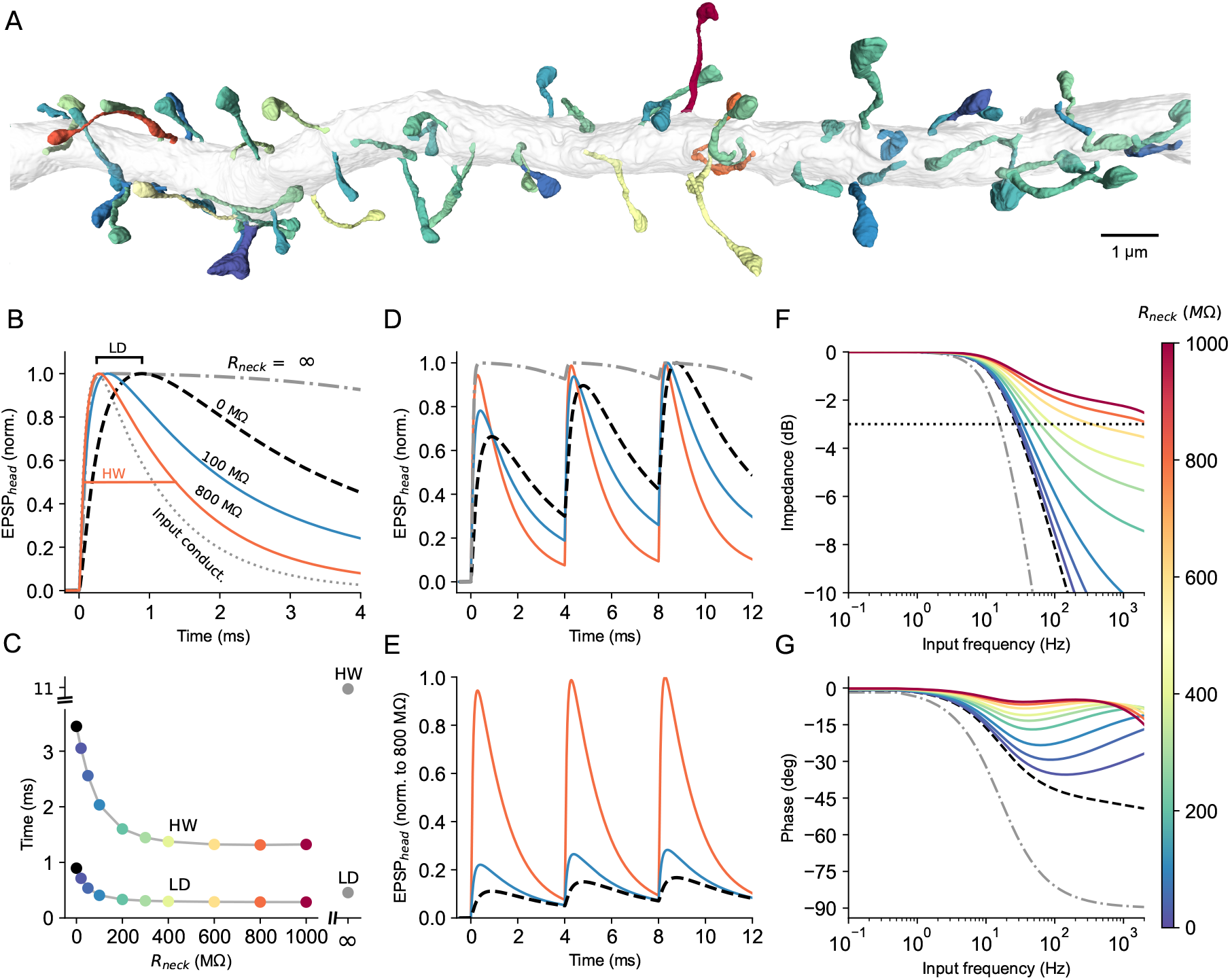
Elevated spine-neck resistance sharpens spine-head EPSPs and supports tracking of high input frequency. **A.** EM-reconstructed dendritic branch (part of the dendrite shown in Figure 1); spines are color-coded by their respective *R*_*neck*_ value, computed using the section-by-section method. Color scale at right. **B**. Normalized EPSPs at the spine-head membrane evoked by a brief excitatory synaptic conductance (dotted trace) for four representative *R*_*neck*_ values. Local delay (LD) is the time difference between the peaks of the input conductance and the EPSP peak (illustrated for *R*_*neck*_ = 0*M*Ω). Half-width (HW) is the EPSP width at 50% of its peak (illustrated in orange for *R*_*neck*_ = 800*M*Ω). **C**. LD and HW versus *R*_*neck*_. Gray circles at right are the LD and HW values for an isolated/isopotential spine (*R*_*neck*_ = ∞). **D**. Normalized spine-head EPSPs in response to three consecutive inputs for the four cases shown in B. **E**. As in D, but all traces are normalized to the peak of the largest (orange) EPSP (in spine with *R*_*neck*_ = 800*M*Ω). The EPSPs in the isolated-spine case are omitted for clarity. **F-G**. Spine-head membrane impedance magnitude (F) and phase (G) as a function of sinusoidal input frequency applied at the spine head for different *R*_*neck*_ values. Dotted horizontal line in F marks the cutoff frequency at −3 dB. The dendritic model consists of a 25*λ* long cylindrical passive cable with a 1*µm* diameter and a single spine in its middle. *C*_*m*_ = 1*µF/cm*^2^; *R*_*m*_ = 10, 000Ω*cm*^2^ (resulting in *τ*_*m*_ = 10*ms*), and *R*_*i*_ = 100Ω*cm*. Each reconstructed spine was represented by an isopotential head and “equivalent cylindrical neck” that preserves both *R*_*neck*_, calculated using the section-by-section method, and the total neck membrane area (see **Methods**). Synaptic parameters: *E*_*AMPA*_ = 0*mV*; *g*_*max*_ = 0.4*nS* with *τ*_*rise*_ = 0.1*ms* and *τ*_*decay*_ = 1*ms*, and *E*_*rest*_ = *™*70*mV* (see **Methods**).

Increasing spine-neck resistance within the biologically feasible range of 100*M*Ω *™* 1*G*Ω progressively sharpens the EPSP at the spine head (Figure 3B). Specifically, the EPSP rises faster (smaller local delay, LD) and becomes narrower (smaller half-width, HW). This effect is quantified in Figure 3C, showing that when *R*_*neck*_ increases between 0 − 400*M*Ω, both LD and

HW are reduced by a factor of ∼3; LD reduces from ∼1*ms* to ∼0.3*ms*, and the HW from ∼4*ms* to ∼1.3*ms*. Noteworthy is that the time-course of the EPSP in the limiting case of an isolated spine (where *R*_*neck*_ = ∞) is significantly longer with LD = 0.5 ms and HW = 11 ms (grey dash-dotted line in Figure 3B) compared to the cases where the spine is coupled via *R*_*neck*_ to the dendrite. The strong effect of *R*_*neck*_ on the dynamics of spinous EPSP is elaborated in Figure 4, and explained mathematically in **Appendix B**. Together, these results identify *R*_*neck*_ as a key determinant of the integration time-window at the individual spine head, with larger *R*_*neck*_ producing briefer EPSPs.

**Figure 4.**
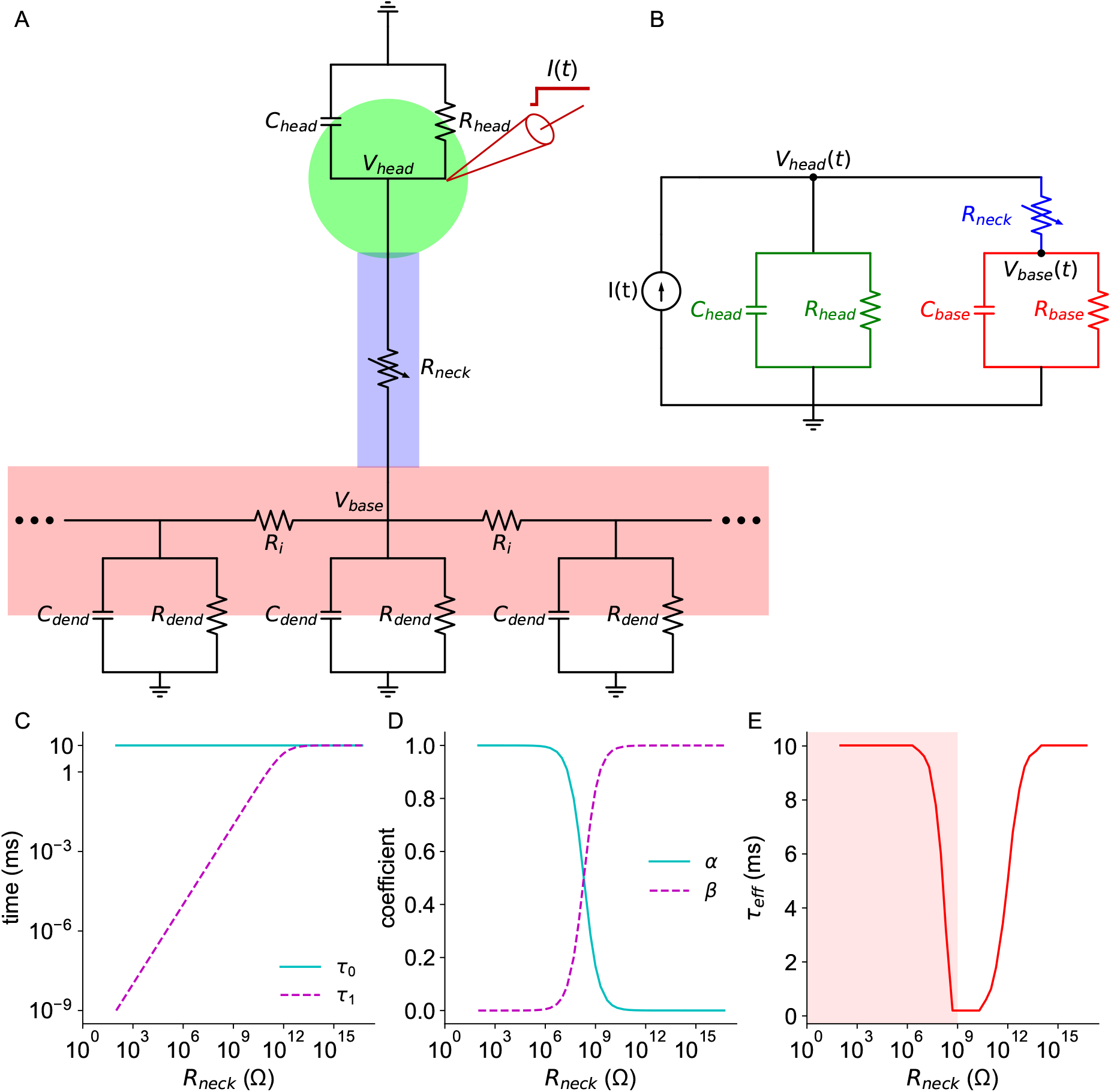
Origin of the ultra-fast dynamics at the spine head membrane. **A.** Equivalent circuit of the spine head compartment (green) coupled, via the spine neck resistance (blue), to the dendritic compartments (red). The respective capacitances and resistances are marked for each compartment. Step current, *I*(*t*), is injected into the spine head (schematic electrode); the resulting voltage response at the spine head and base are denoted by *V*_*head*_ and *V*_*base*_, respectively. **B**. Reduced two-compartment model of the circuit shown in A. The voltage response in this circuit is governed by a slow (*τ*_0_) and a fast (*τ*_1_) time constant (Equation 6). **C-E**. Analysis of the reduced circuit shown in B. **C**. The equalizing time constant, *τ*_1_ (dashed magenta), and the slow time constant, *τ*_0_ (cyan), as a function of *R*_*neck*_. **D**. The coefficients *α* and *β* that determine the relative contribution of the two exponential components, governed by *τ*_0_ and *τ*_1_, respectively (Equations 7 and 8), as a function of *R*_*neck*_. **E**. Effective time constant, *τ*_*eff*_, as a function of *R*_*neck*_. *τ*_*eff*_ is the time at which *V*_*head*_ reaches 63% of its steady-state value following step current input (see Appendix B). The shaded area depicts the biologically feasible range of *R*_*neck*_. Model parameters: *C*_*m*_ = 1*µF/cm*^2^; *R*_*m*_ = 10, 000Ω*cm*^2^ (yielding *τ*_*m*_ = 10*ms*) for both the spine and dendrite. The spine head membrane is 1*µm*^2^, yielding *R*_*head*_ = 1*T* Ω and *C*_*head*_ = 0.01*pF*; *R*_*base*_ and *C*_*base*_ = 200*M*Ω and 0.05*nF*, respectively.

The sharpening of the EPSP as a function of *R*_*neck*_ implies that the temporal summation at the spine head membrane is strongly influenced by *R*_*neck*_. Indeed, repeated activation of the synapses at 250*Hz* results in negligible temporal summation for large *R*_*neck*_ (e.g., 800*M*Ω; orange trace) but pronounced temporal summation for small *R*_*neck*_ (e.g., 100*M*Ω; blue trace) and even more so when the synapses contact directly the dendritic cable (*R*_*neck*_ = 0*M*Ω; dashed black trace) (Figure 3D). Temporal summation is most evident in the electrically isolated spine (*R*_*neck*_ = ∞; grey dash-dotted trace), consistent with the substantially prolonged EPSP time course in this case. As expected from previous theoretical studies (Segev and Rall, 1988), the *R*_*neck*_-dependent sharpening of the spinous EPSPs is accompanied by an increase in EPSP amplitude (Figure 3E). Together, Figures 3D and 3E demonstrate that, within the biological-range of *R*_*neck*_, AMPA-like spinous EPSPs are expected to be both brief and large, optimizing them for precise local computation (see also, Gulledge et al. (2012)).

Enhanced spine-head EPSP dynamics with increasing *R*_*neck*_ result in improvement in tracking high-frequency input modulations at the spine head. This is quantified in Figure 3F, where the impedance (in dB) as a function of sinusoidal input frequency at the spine head is depicted. In the limiting cases of an isolated spine (grey dash-dotted line) and a spine lacking a neck (black dashed line), the −3 dB cutoff frequencies are relatively low (15*Hz* and 30*Hz*, respectively). In contrast, for intermediate-to-large *R*_*neck*_ values, the cutoff frequency shifts to substantially higher values (e.g., ∼90*Hz* for *R*_*neck*_ = 400*M*Ω, increasing to ∼3*kHz* for *R*_*neck*_ = 1*G*Ω) (Figure 3F). Consistent with this, the phase response shows reduced lag for large *R*_*neck*_ (Figure 3G).

To conclude, elevated *R*_*neck*_ markedly accelerates spine-head EPSP kinetics as compared to a shaft synapse (where *R*_*neck*_ = 0*M*Ω), narrowing the effective integration time-window at the spine head, thereby markedly enhancing the spine’s ability to track high-frequency inputs.

### Origin of the ultra-fast dynamics of spinous EPSPs

To explain the observation that *R*_*neck*_ gives rise to fast voltage dynamics at the spine head, we analytically solve a reduced two-compartment case of a spine connected to a dendritic branch. Figure 4A shows the equivalent circuit model of a dendritic spine connected to a dendritic cable via the spine neck resistance, *R*_*neck*_. This circuit was simplified to a two-compartment model illustrated in Figure 4B. In this lumped model, the voltage response to a current injected at the spine head can be expressed as the sum of two exponential terms, with time constants *τ*_0_ and *τ*_1_ and respective coefficients *α* and *β* (Equations 6 - 8; see full derivation in **Appendix B**.; see also Golowasch et al. (2009)).

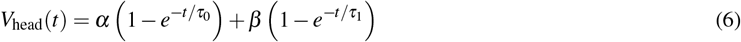

when the specific membrane properties (*C*_*m*_, *R*_*m*_) are identical for both spine and dendrite, *τ*_0_ = *τ*_*m*_ = *C*_*m*_*R*_*m*_ is the slower (membrane) time constant, whereas *τ*_1_ is a faster “equalizing” time constant (Rall, 1969). In this case,

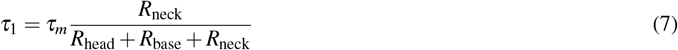

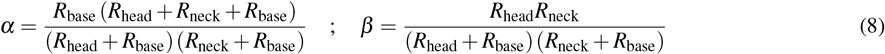

Figure 4C shows that *τ*_1_ varies steeply with *R*_*neck*_, attaining extremely small values over biological values of *R*_*neck*_ and approaching *τ*_0_ at very large *R*_*neck*_. Figure 4D shows the values of the respective *α* and *β* weights; at low *R*_*neck*_, the coefficient for *τ*_0_ dominates (*α ≈* 1, *β≈* 0), whereas at very large *R*_*neck*_ the coefficient for *τ*_1_ dominates (*β ≈* 1, *α≈* 0). In this case, the fast time constant, *τ*_1_, dominates the voltage dynamics at the spine head membrane.

To quantify the effect of *R*_*neck*_ on the time course of the voltage at the spine head, *V*_*head*_(*t*), we computed the effective time constant, *τ*_*eff*_, defined as the time required for *V*_*head*_(*t*) to reach 63% of its steady-state value following a step current applied at the spine head (Figure 4E). *τ*_*eff*_ decreases sharply with increasing *R*_*neck*_ over the low-to-intermediate range of *R*_*neck*_, reaching a minimum of ∼200*µs* for the parameters used at *R*_*neck*_ *≈*1*G*Ω. This minimum occurs in the regime where *τ*_1_ is very small, and the respective coefficient *β* is substantial. For very large (unrealistic) *R*_*neck*_, *τ*_1_ converges to *τ*_0_ (Figure 4C), and the voltage response at the spine head becomes slow. Simulation results for *τ*_*eff*_ as a function of *R*_*neck*_ in the full circuit of Figure 4A can be found in **Appendix B**.

In conclusion, an analytic solution for the two-compartment model representing spine head compartment connected, via *R*_*neck*_, to a dendrite compartment provides an explanation for the ultra-fast voltage dynamics in the spine head found in our simulations. Intuitively, for the biological range of *R*_*neck*_ (100*M*Ω *™*1*G*Ω), the dendritic compartment serves as a substantial current “sink” for the much smaller spine head compartment, yielding a fast equalizing time constant, *τ*_1_, that enhances the voltage response at the spine head. This mechanism explains the narrowing of spinous EPSPs with an increase in *R*_*neck*_, shown in Figures 3B and 3C.

### Impact of spine density on the tracking of fast spinous inputs

Spine density varies markedly among dendritic branches (Figure 5A); this is expected to influence the impedance load (i.e., current sink) that the dendritic branch imposes on individual spines. We next examined how this “spine environment” affects the EPSP time course at the spine head and, consequently, the ability of the spine to track high-frequency synaptic inputs.

**Figure 5.**
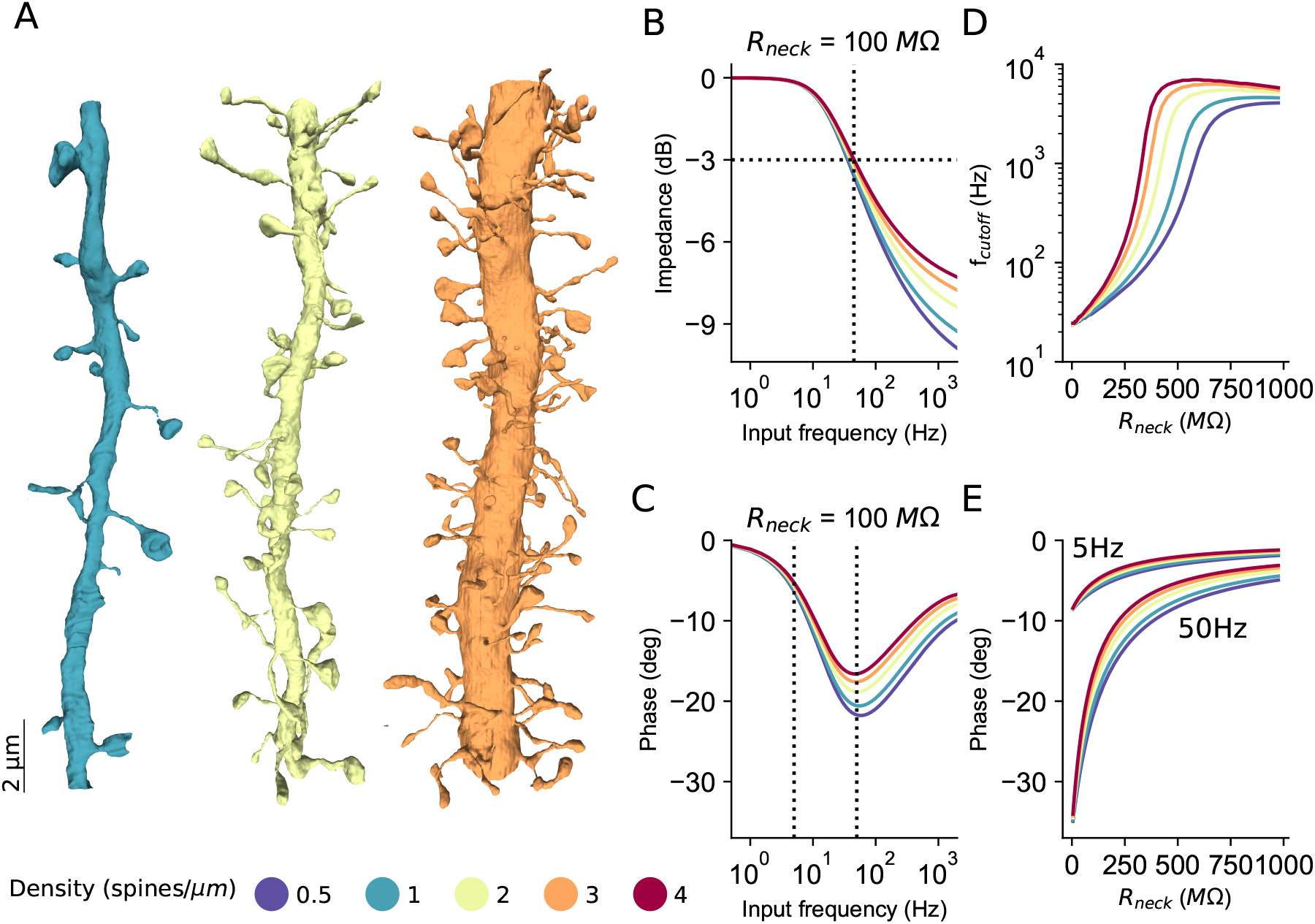
Input tracking in spines is enhanced with an increase in spine density. **A.** Three representative dendritic branches with different spine densities (color-coded by circles at bottom). **B, C**. Membrane impedance and phase shift, respectively, at the spine-head membrane as functions of input frequency for different spine densities. All modeled spines have *R*_*neck*_ = 100*M*Ω. Horizontal dashed line in B marks the cutoff frequency at −3 dB. Vertical dashed line marks 45*Hz*; the intersection point between the two lines depicts the cutoff frequency for a density of 4 spines/*µm* (red curve). The vertical dashed lines in C denote 5*Hz* and 50*Hz* input frequencies – further analyzed in E. **D**. Cutoff frequency as a function of *R*_*neck*_ for different spine densities. Increasing spine density shifts the cutoff to higher frequencies. **E**. Phase shift at 5*Hz* and 50*Hz* as a function of *R*_*neck*_ and spine density. Model: near-infinite cable (as in Figure 4) with the specified spine density. Spine density was modeled using the F-factor (see **Methods**).

Increasing spine density markedly enhances the tracking of high-frequency inputs at the input spine, as reflected by larger cutoff frequencies (−3 dB) and reduced phase lag with an increase in spine density (Figures 5B-C). This effect results from the shortening of both the local delay (LD) and the half-width (HW) of the spinous EPSP at the input spine with an increase in spine density.

The combined effects of *R*_*neck*_ and spine density on the cutoff frequency and phase shift are summarized in Figures 5D-E. Increasing *R*_*neck*_ shifts the cutoff frequency to higher values while reducing the phase shift. Notably, increasing spine density has a similar effect to increasing the diameter of the parent dendrite: both accelerate the EPSP at the input spine and enhance its ability to follow rapid input modulations.

### Strong and temporally precise inhibitory gating in dually-innervated spines (DiSs)

Dually innervated spines (DiSs), which receive both excitatory (E) and inhibitory (I) inputs on the same spine, have been described throughout neocortex and hippocampus (Knott et al., 2002; Kubota et al., 2007; Chen et al., 2012; Kleinjan et al., 2023; Gemin et al., 2021). Because the local voltage dynamics in DiSs are dominated by the fast equalizing time constant *τ*_1_ (Figure 4), we expect that spine inhibition will exert a strong and temporally precise gating effect on the co-located excitatory input.

We quantify the effect of inhibition in DiSs by defining the inhibitory gate as the fractional reduction in EPSP peak (or area under the EPSP or *Ca*^2+^) caused by inhibition,

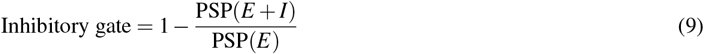

where PSP(E+I) is the PSP parameters measured when both E and I are active, and PSP(E) is the respective case when only E is active. The inhibitory gate equals 0 when inhibition has no effect and 1 when inhibition completely vetoes the PSP-related parameters.

Figure 6 illustrates that, under synchronous activation of E and I in DiSs, “silent” inhibition (whose battery equals the resting potential) very effectively suppresses the spinous EPSP, and that this effect becomes markedly stronger as *R*_*neck*_ increases. Figure 6A presents an EM reconstruction of a DiS; the inset shows the modeled E and I synaptic conductances, and the corresponding equivalent circuit is shown in Figure 6B. An example of the inhibitory effect on the EPSP for *R*_*neck*_ = 100*M*Ω (blue) and 800*M*Ω (orange) is shown in Figure 6C for PSPs recorded at the spine head, and in Figure 6D for PSPs recorded at the spine base. Note that, surprisingly, for the synaptic parameters used here, the percentage reduction of the EPSP peak is larger at the dendritic base (and thus at the soma) than at the spine head, even though both E and I synapses are located at the spine head. In other words, the relative difference in peak PSP with and without inhibition is greater at the spine base (Figure 6D) than at the spine head (Figure 6C) for both *R*_*neck*_ values examined.

**Figure 6.**
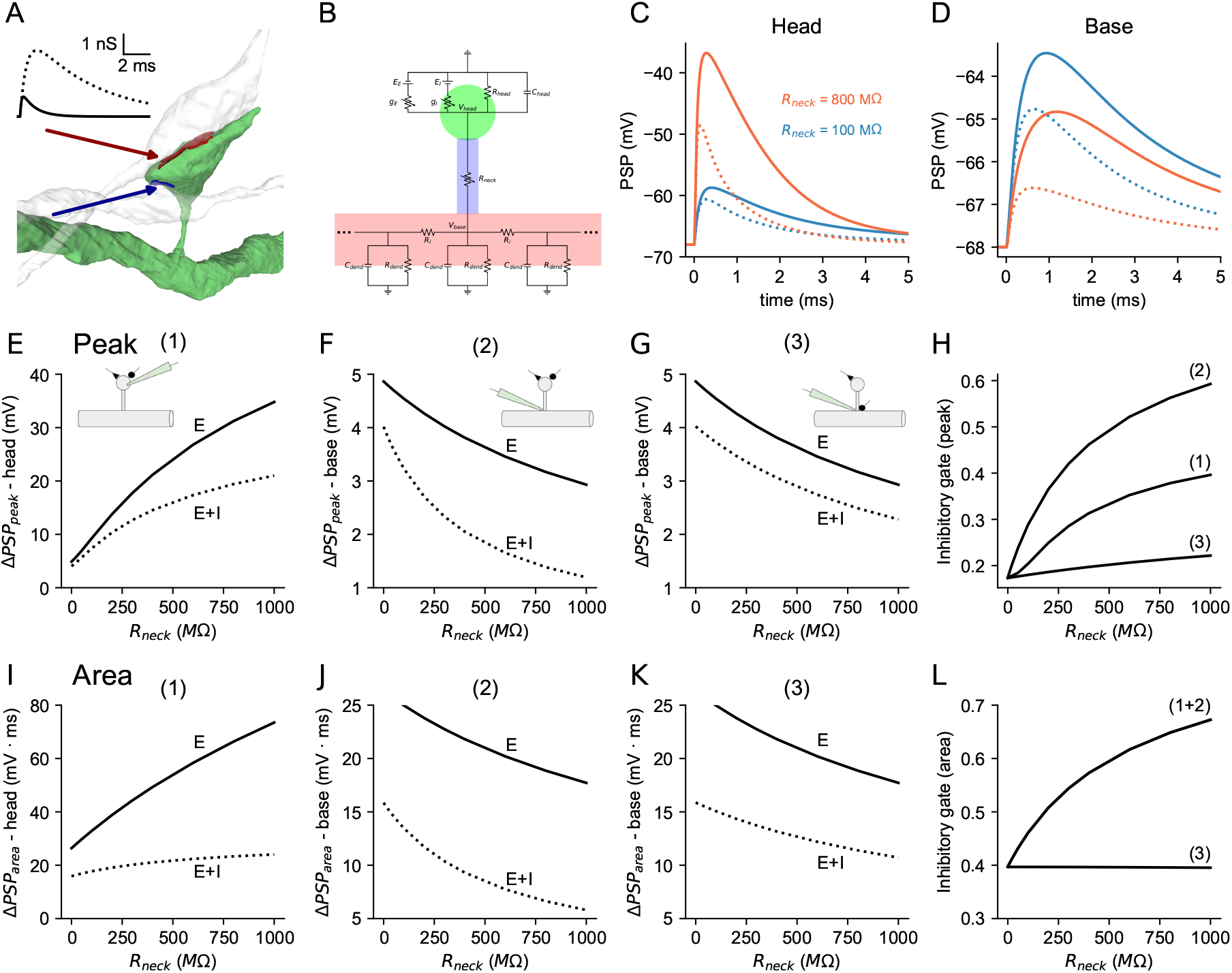
Degree of inhibitory gating in dually innervated dendritic spines depends on neck resistance. **A.** An EM-reconstruction of dually-innervated spines, DiS. The two synapses are shown in red and blue. The inset shows the amplitude and time-course of the excitatory (continuous line) and inhibitory (dotted line) conductances. **B**. An equivalent circuit model for a DiS; the dendritic branch is shown in red. **C**. PSPs at the spine head with inhibition (dotted line, E+I) and without inhibition (continuous line, E) for *R*_*neck*_ = 100*M*Ω (blue) and 800*M*Ω (orange). **D**. As in C, but the resultant PSPs are at the spine base. **E**. PSP peaks at the spine head for E-synapse alone and for E+I synapses, as a function of *R*_*neck*_. **F**. As in E, but at the spine base. **G**. PSP peaks at the spine base when E-synapse is at the spine head, and I-synapse is at the spine base. **H**. Inhibitory gate (Equation 9), for the PSP peak. **I-L**. As in E-H, but for the PSP area. Model parameters: *g*_*AMPA*_: *τ*_*rise*_ = 0.1*ms, τ*_*decay*_ = 1*ms, g*_*max*_ = 1*nS. g*_*GABA*_: *τ*_*rise*_ = 0.5*ms, τ*_*decay*_ = 5*ms, g*_*max*_ = 3.3*nS. E*_*AMPA*_ = 0*mV*; *E*_*GABA*_ = *E*_*pass*_ = −68*mV*.

Figure 6E shows the effect of inhibition on the EPSP peak at the spine head as a function of *R*_*neck*_; the corresponding PSPs’ peak at the spine base is shown in Figure 6F. As *R*_*neck*_ increases, the PSP at the spine head increases for both E and E+I conditions (Figure 6E), whereas the respective PSPs at the spine base decrease (Figure 6F). A control configuration, in which excitation is at the spine head and inhibition is at the spine base, yields substantially weaker inhibitory gating (Figure 6G). The same set of comparisons was repeated using PSP area (time integral) rather than PSP peak, providing a time-integral-based view of inhibitory gating in DiSs gating (Figures 6I-L).

The dependence of inhibitory gating on *R*_*neck*_ is summarized in Figures 6H and 6L. *R*_*neck*_ acts as a “knob” that strongly modulates the strength of inhibitory gating. When inhibitory gating is evaluated at the PSP peak (Figure 6H), it reaches ∼0.4 at the spine head (case 1) and∼ 0.6 at the spine base (case 2) for *R*_*neck*_ = 1000*M*Ω. In contrast, inhibitory gating is much weaker (∼ 0.15) when excitation occurs at the spine head and inhibition is applied at the spine base (case 3). Note that for the PSP peak (Figure 6H), although inhibition is applied at the spine head, its gating effect is larger at the base than at the head. This surprising asymmetry arises from the relatively slow inhibitory conductance used in this model, which partially “misses” the earlier EPSP peak at the head but effectively “catches up” with and dampens the delayed EPSP peak at the base. For inhibitory gating over the PSP area (Figure 6L), the gating at the head and base are identical as a function of *R*_*neck*_ (cases 1 and 2), reaching ∼0.7 for *R*_*neck*_ = 1000*M*Ω; it remains weaker, ∼0.4, when inhibition is located at the spine base (Figure 6L, case 3). Additional cases, including different E and I time courses and inhibitory gating of EPSP area rather than peak, are presented in Supplementary Figures S2-S7.

Taken together, these results identify *R*_*neck*_ as a structural control mechanism (a “knob”) that enables highly effective, *R*_*neck*_-dependent inhibitory gating of co-localized excitation (Figures 6H, and 6L). Possible implications of this result for information routing and continual learning are considered in the **Discussion**.

We next study the dependence of the inhibitory gating as a function of time difference (Δ*t*) between E and I activation. Figure 7A shows six examples of normalized PSPs at the spine head; the top pair corresponds to Δ*t* = *™*1*ms* (I before E), the middle pair to Δ*t* = 0*ms* (I coincident with E), and the bottom pair to Δ*t* = +1*ms* (I after E), for the case of *R*_*neck*_ = 100*M*Ω (blue) and 800*M*Ω (orange). The EPSPs are shown by the continuous black line and the PSPs (with inhibition) by the dotted line; the difference between the two cases is highlighted by the shaded area. The reduction in the EPSP area following inhibition increases with *R*_*neck*_ for all three cases shown, indicating that the inhibitory gating is more potent with increasing *R*_*neck*_ for all timing conditions.

**Figure 7.**
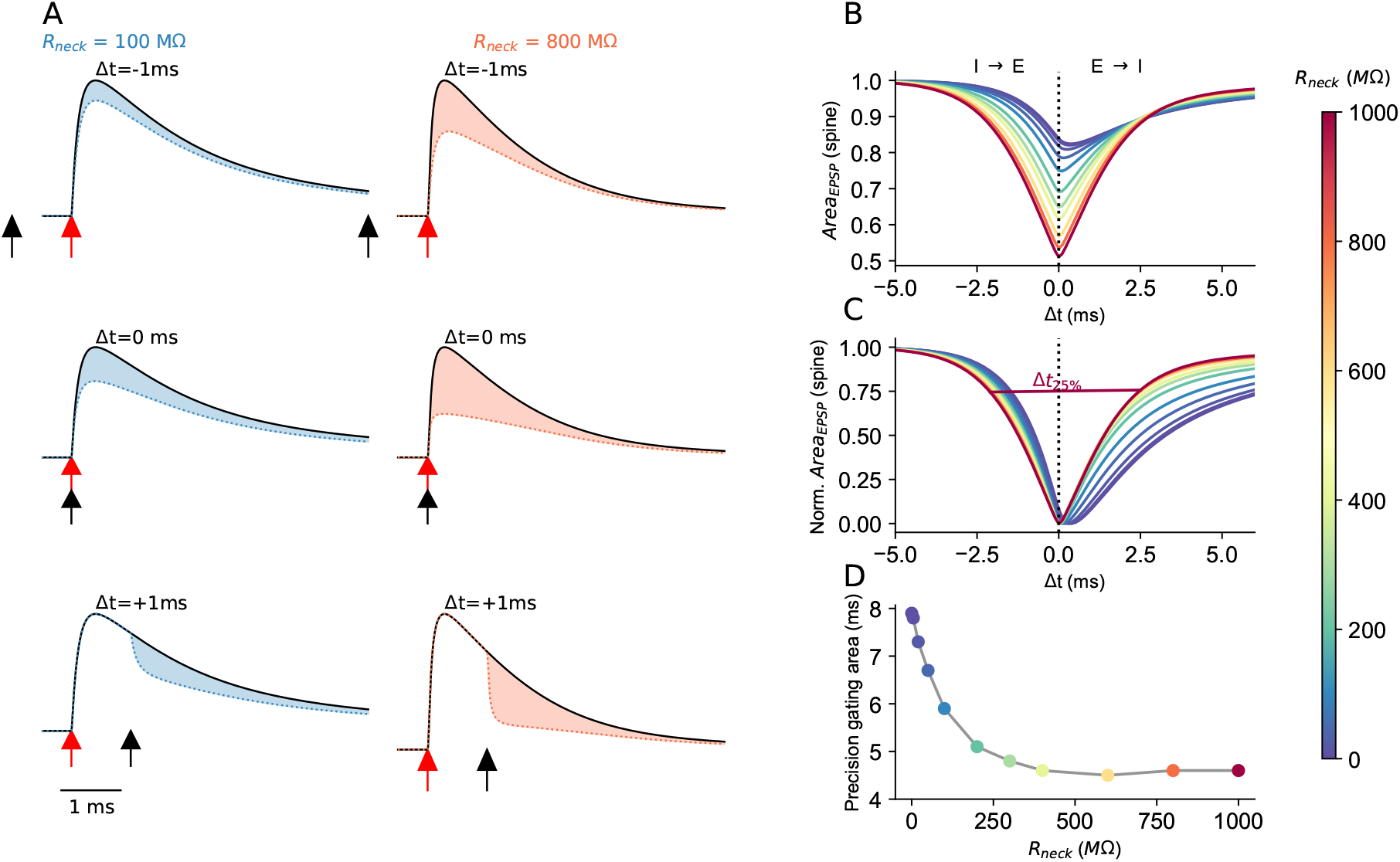
Temporal precision of spinous inhibition depends on spine neck resistance. **A.** Normalized PSPs at the spine head for *R*_*neck*_ = 100*M*Ω (blue) and 800*M*Ω (orange). Shown are three cases of Δ*t* - the time difference between I activation (black arrow) and E activation (red arrow). **B**. Normalized PSP area as a function of Δ*t* for various *R*_*neck*_ values (color code at right). **C**. As in B, but all curves are normalized to range between 0-1; the temporal precision gating of I over E (Δ*t*_25%_) is the width of the curves at 25% reduction of the PSPs area. **D**. Δ*t*_25%_ of the EPSP area as a function of *R*_*neck*_. Model is as in Figure 6, but with faster *g*_*GABA*_ kinetics that are identical to those of *g*_*AMPA*_.

Figure 7B shows that the potency of inhibitory gating measured on the EPSP time-integral (area) at the spine head varies systematically with *R*_*neck*_ (color-coded) as a function of Δ*t*; it is indeed increasingly stronger with an increase in *R*_*neck*_. To quantify the temporal precision with which inhibition gates the PSP time integral, we scaled the curves in Figure 7B so that they all span between 0 and 1 (Figure 7C). We define the temporal precision of inhibitory gating as the width of the curve at 25% reduction in PSP area (Δ*t*_25%_) showing that it decreases monotonically with *R*_*neck*_ (Figure 7D); the temporal precision window of inhibitory gating becomes roughly twice as narrow as *R*_*neck*_ increases from 0 *™* 1000*M*Ω, implying that faster spine-head dynamics compress the effective inhibitory gating to a tighter temporal coincidence window between excitation and inhibition. Note that the Δ*t*-dependence on *R*_*neck*_ is asymmetric around Δ*t* = 0 (Figures 7B, C); the inhibitory gating is broader for Δ*t >* 0 compared to Δ*t <* 0, consistent with the slower decay of the EPSP versus its rise time. Additional cases for E/I temporal interactions in DiSs are provided in Supplementary Figures S5-S7. Together, Figures 6 and 7 demonstrate that DiSs provide an *R*_*neck*_-dependent structural mechanism for both highly effective and temporally precise inhibitory gating.

### *Ca*^2+^ concentration, inhibitory gating, and E/I timing in nonlinear spines

Experimental work has shown that spine heads contain active conductances and that synaptic activation can recruit substantial NMDAR-mediated *Ca*^2+^ influx, shaping local electrical and biochemical signals (Yuste et al., 1999; Mainen et al., 1999; Sabatini et al., 2002; Noguchi et al., 2005; Ngo-Anh et al., 2005; Araya et al., 2007; Miyazaki and Ross, 2017). Modeling studies further showed that spine-head EPSP amplification depends nonlinearly on spine-neck resistance, *R*_*neck*_ (Segev and Rall, 1988; Harnett et al., 2012; Doron et al., 2017). Here we examine how an excitatory synapse with AMPA + NMDA components interacts with a co-localized inhibitory synapse in DiSs, and how this interaction shapes *Ca*^2+^ elevation, inhibitory gating, and E/I timing dependence.

In the absence of inhibition (Figure 8A, solid lines), increasing *R*_*neck*_ nonlinearly boosts spine-head depolarization, thereby enhancing NMDAR recruitment during the EPSP and elevating the resulting *Ca*^2+^ transient (Figure 8B). When inhibition is coactive (Figure 8A, dotted traces), spine-head depolarization is reduced in an *R*_*neck*_-dependent manner, consistent with a stronger local impact of inhibitory conductance at larger *R*_*neck*_. As *R*_*neck*_ increases, inhibition more effectively gates NMDAR-driven *Ca*^2+^ entry, flattening the *Ca*^2+^ peak across a broad range of *R*_*neck*_ values (Figure 8C).

**Figure 8.**
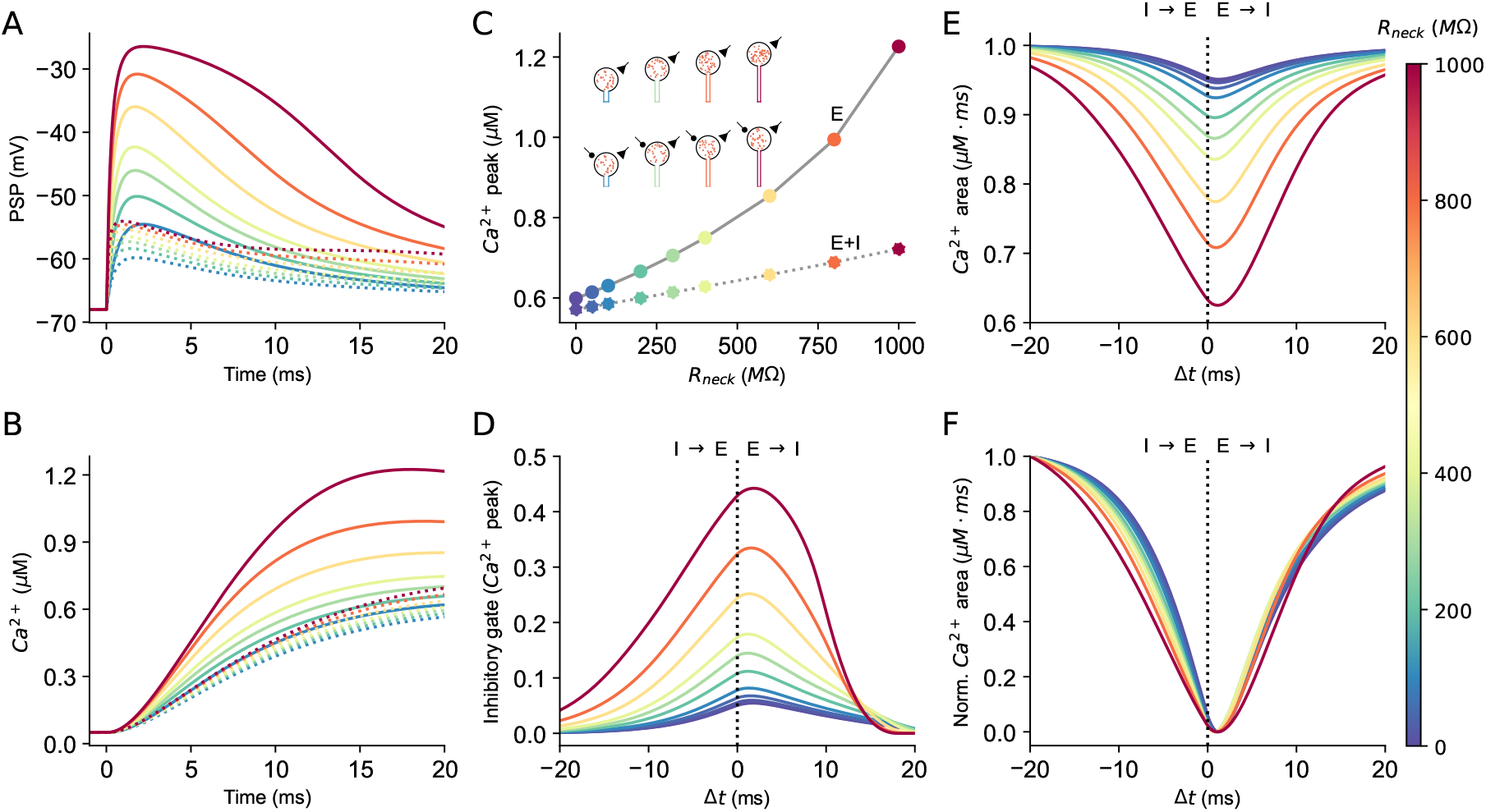
Spine neck resistance amplifies NMDA-dependent *Ca*^2+^ signals and augments inhibitory gating. **A.** Spine head PSPs in response to AMPA + NMDA-based E-synapse (continuous lines) and simultaneous E+I synapses (respective dotted lines) for a range of *R*_*neck*_ values (color code at right). **B**. Corresponding NMDA-mediated *Ca*^2+^ concentration for the cases shown in A. **C**. Peak NMDA-mediated spine-head *Ca*^2+^ concentration as a function of *R*_*neck*_ for E alone and simultaneous E+I. The inset shows schematics of 4 exemplar spines (*R*_*neck*_ = 100, 300, 800, 1000*M*Ω) without (top) and with inhibition (bottom); of dots/spine is proportional to the *Ca*^2+^ concentration at the spine head. **D**. Inhibitory gate (Equation 9) for the peak of *Ca*^2+^ concentration as a function of Δ*t* for various *R*_*neck*_ values. **E**. Normalized area of *Ca*^2+^ concentration as a function of Δ*t* for various *R*_*neck*_ values. **F**. As in E, but all curves are normalized to range between 0 and 1. Model parameters as in Figure 6, here with *g*_*NMDA*_: *τ*_*rise*_ = 3*ms, τ*_*decay*_ = 30*ms, g*_*max*_ = 2*nS*; *E*_*NMDA*_ = 0*mV*.

To quantify timing specificity, we computed the inhibitory gate for the *Ca*^2+^ peak (Equation 9) as a function of time delay Δ*t* between E and I activation (Figure 8D). Gating strength increases markedly with *R*_*neck*_: for example, peak gating is small (∼0.05) for *R*_*neck*_ = 100*M*Ω and ∼10 larger for *R*_*neck*_ = 1000*M*Ω. An analogous analysis of the *Ca*^2+^ time integral (Figures 8E, F) shows complementary gating of total *Ca*^2+^ concentration. Because NMDAR kinetics are slow, the temporal inhibitory gating window is less sensitive to *R*_*neck*_ than in the AMPA-only case (compare Figure 7C to Figure 8F).

In summary, increasing *R*_*neck*_ nonlinearly amplifies NMDAR-dependent spine-head *Ca*^2+^ signals. In DiSs, coactive inhibition gates both the *Ca*^2+^ peak and its time integral as a function of *R*_*neck*_, effectively stabilizing spine *Ca*^2+^ across a broad range of *R*_*neck*_ values.

## Discussion

This work combines dense EM reconstructions with analytic theory and biophysical modeling to ask how nano-scale features of dendritic spines shape the temporal processing of spinous synaptic inputs and the strength and time-window of inhibitory gating in cortical pyramidal neurons. By systematically analyzing thousands of EM-reconstructed spines, we show that spine necks often exhibit repeated local constrictions (Figure 1) that substantially increase spine-neck resistance (*R*_*neck*_, Figure 2). Incorporating these realistic neck geometries into compartmental models of spinous dendrites reveals that *R*_*neck*_ is a powerful “timing knob”. It introduces a fast equalizing time-constant to the spine-dendrite system, thereby shortening the effective integration time window for spinous EPSPs. This in turn improves the tracking of fast synaptic modulations (Figures 3 and 4), and strongly regulates the potency and temporal precision of inhibition in regulating the local voltage and *Ca*^2+^ signaling in dually innervated spines (DiSs; Figures 6 - 8). Together with our simulations of EM-reconstructed spiny dendritic branches (Figure 5), these results position dendritic spines as key temporal processors of their synaptic inputs, and modulators of inhibitory gates in DiSs, thus linking spine ultrastructure to the computational repertoire and synaptic plasticity of cortical pyramidal neurons.

### EM-level reconstruction reveals repeated spine-neck narrowing, especially in long-neck spines

EM-based analysis of thousands of dendritic spines revealed a geometric motif – repeated local narrowing along the spine neck. It was shown previously that spines show changes in neck diameter along their length (Tamada et al., 2020; Benavides-Piccione et al., 2025); here, we found that spine necks frequently exhibit highly constricted segments, having local diameters of only a few tens of nanometers along the spine neck (Figure 1). To quantify the impact of such constrictions on *R*_*neck*_, we compared the traditional “average” diameter method, which collapses the neck into a single “average” cylinder, with our new “section-by-section” method that sums the neck resistances from a series of short EM-defined segments (Figure 2A). We showed analytically that the latter approach yields substantially higher *R*_*neck*_ than the former. In our dataset of 2,074 spines, the average-diameter method systematically underestimates *R*_*neck*_ by ∼50%, with the discrepancy between the two methods increasing as neck geometry becomes more irregular (Figures 2B-C).

This repeated narrowing of the spine neck is not randomly distributed; spines with long necks are much more likely to contain very thin constrictions, leading to markedly elevated *R*_*neck*_ in these spines. Indeed, the MAD index we have used to assess diameter irregularity is positively correlated with the neck length and with *R*_*neck*_ (Figures 1J and 2E). Thus, long-neck spines are not simply a stretch of short-neck spines; they are enriched in nano-scale bottlenecks that strongly shape their electrical properties. Only EM-level reconstructions with segment-wise biophysical analysis reveal this landscape of spine resistance, and hence the extent to which spine morphology controls the strength and timing of synaptic potentials and *Ca*^2+^ signaling, and potency of inhibitory gating at the single-spine level.

### Spine necks as dynamic temporal filters

A central result of our study is that increasing *R*_*neck*_ not only increases the local voltage at the spine head and attenuates synaptic potential from the spine head to the parent dendrite and soma, as originally demonstrated theoretically by Rall (1974) and validated experimentally (Bloodgood and Sabatini, 2005), but also accelerates local voltage dynamics. Analytic treatment of a spine-on-cable model (Figure 4 and **Appendix B**.) shows that neck-resistance introduces an additional equalizing time constant (*τ*_1_) that can become much shorter than the membrane time constant (*τ*_*m*_). We show that for a range of biologically-realistic *R*_*neck*_ values (100 *™* 1000*M*Ω), the neck-dominated fast equalizing time constant sharpens the EPSPs (and IPSPs) at the spine head by up to ∼3-fold (Figure 3C). As a result, spines with large *R*_*neck*_ behave as ultra-fast local filters that can faithfully follow synaptic fluctuations in the range of 100-Hertz; a regime in which synaptic inputs impinging directly on the stem dendrite would exhibit strong low-pass filtering (Figure 3).

These findings extend classic cable-theory predictions for dendrites and spines by grounding them in realistic connectomic morphologies. Prior modeling and experimental studies have emphasized that spines are electrical and biochemical micro-compartments whose resistance and small head volume shape local depolarization, encourage local nonlinearities and enhance *Ca*^2+^ entry, and plasticity processes (Segev and Rall, 1988; Koch and Poggio, 1983; Yuste et al., 2000). Our analysis further shows that once the full tortuosity and constriction of EM-resolved necks are taken into account, their impact on temporal dynamics is much stronger than expected from cylindrical spine necks with “average” diameter. In this sense, the biophysical consequences of *R*_*neck*_, in particular at the time domain, may have been systematically underestimated in previous studies.

It is important to note that the experimental value of *R*_*neck*_ is still unsettled: reported estimates span a wide range, from low values of ∼30*M*Ω in “mushroom-shaped” spines (Popovic et al., 2015; Weng et al., 2025), up to several hundred *M*Ω in apical spines (Harnett et al., 2012; Jayant et al., 2017), with recent GEVI-based measurements reaching 530*M*Ω (Cornejo et al., 2022). Because direct measurements are difficult, many studies infer *R*_*neck*_ from morphology and assume specific axial resistivity (Rall, 1974; Harris and Stevens, 1989; Svoboda et al., 1996; Gemin et al., 2021). Using our EM-based “section-by-section method” and assuming *R*_*i*_ = 150Ω*cm* as is common in the literature (Tønnesen et al., 2014; Cartailler et al., 2018; Tamada et al., 2020), we obtained *R*_*neck*_ values ranging between 2*M*Ω and 920*M*Ω for our spine population. However, intracellular organelles such as the spine apparatus can occlude 7 - 27% of the neck lumen (Wilson et al., 1983; Harris and Stevens, 1989), and their presence correlates with neck geometry (Ofer et al., 2021), likely increasing effective *R*_*neck*_ beyond our estimates. For example, assuming *R*_*i*_ = 300Ω*cm* as in e.g., (Gemin et al., 2021), would double all *R*_*neck*_ values reported in the present study. We note that we detected small segments with a very thin spine neck (5*nm*); in addition to the large contribution of these segments to *R*_*neck*_, they may also restrict ion (e.g., *Ca*^2+^) diffusion along such ‘bottlenecks’ (Cartailler et al., 2018; Lagache et al., 2019).

### Trade-off between local precision and branch-level integration

By embedding EM-derived spine density into dendritic cable models (Figure 5), we demonstrate that there is a trade-off between spine density and *R*_*neck*_ regarding local temporal precision and signal propagation along the dendritic cable. EPSPs in the input spine are accelerated with an increase in both spine density and *R*_*neck*_, thereby enhancing the tracking of fast spinous inputs. At the same time, increasing spine density slows signal propagation along the dendritic branch, because each spine acts as a capacitive and resistive load that “sinks” axial current flowing along the parent dendrite. This trade-off suggests a structural mechanism for differentiating dendritic segments specialized for precise local coincidence detection versus those optimized for broader spatial integration. Densely spiny branches covered with spines with large *R*_*neck*_ may support temporally precise, locally gated computations. In such branches, local computations/gating operate on sub-millisecond scales, sensitive to fine-grained timing of sensory inputs. Smoother dendritic branches, however, with low *R*_*neck*_ spines, may be better suited for fast distribution and summing inputs over larger dendritic domains. This possibility will be further explored in future studies.

### Dually innervated spines as nano-meter inhibitory gates

Perhaps the most striking computational consequence of high *R*_*neck*_ emerges in dually innervated spines (DiSs), of which several hundreds of them are hosted by each PN (Kubota et al., 2007; Chiu et al., 2013; Boivin and Nedivi, 2018; Bloss et al., 2016; Tremblay et al., 2016; Kubota et al., 2016; Chen et al., 2012; Gemin et al., 2021; Tullis and Bayer, 2024; Müllner et al., 2015). Because the excitatory and inhibitory synapses share the same tiny head compartment, inhibition can gate excitation with exceptional potency (Koch and Poggio, 1983; Gemin et al., 2021). We quantified this effect by defining an “inhibitory gate” index based on the reduction of EPSP peak, time-integral, or *Ca*^2+^ concentration due to the activation of co-located inhibition, and introduced a temporal precision measure to the width of the E-I interaction window. We found that for moderate *R*_*neck*_ values (ranging between 100 *™*400*M*Ω), spinous inhibition achieves very strong and temporally precise control over the co-located EPSP: a narrow band of E-I timing around Δ*t ≈* 0*ms* produces large suppression of the respective EPSP as well as *Ca*^2+^ signals by about 60-70%; this interaction window sharpens as *R*_*neck*_ increases (Figures 6 - 8).

These results suggest that DiSs function as ultra-local, nanoscopic “veto gates” that can switch individual excitatory inputs on or off with millisecond precision. Importantly, this gating is highly sensitive to the nano-geometry of the neck and to the exact timing of inhibitory recruitment, providing a direct mechanistic interpretation for experimental observations that inhibitory synapses on spines are dynamically formed, removed, and repositioned during learning and experience (Jasinska et al., 2010; Caroni et al., 2014; Knott et al., 2002). In circuit terms, populations of DiSs could implement a rich palette of context- and time-dependent filters, selectively permitting (or blocking) specific afferent pathways depending on inhibitory drive and spine state and respective *R*_*neck*_ values.

### Limitations and future directions: toward a “biophysics of connectomics”

Our modeling used passive dendritic cables combined with a limited set of active conductances on the spine membrane to isolate the impact of *R*_*neck*_ and spine density on temporal processing in individual spines. Real cortical neurons and spines have richer active properties, heterogeneous channel density, ongoing synaptic activity, and glial interactions, and our EM data cover only specific layers and cell types. Moreover, we examined fixed morphologies, while *in vivo*, both spine geometry and synaptic contacts, in particular inhibitory ones, are dynamic.

A potential concern is whether the spine-neck constrictions observed here could arise from EM-related fixation or reconstruction artifacts. However, this is difficult to reconcile with the dense packing of the neuropil, where nonspecific tissue shrinkage would be expected to affect surrounding structures similarly rather than producing highly localized narrowing. In addition, a recent study using another method (cryo fixation) also found such spine-neck restrictions (Tamada et al., 2020), and in our visual inspection of publicly available light-microscopy-based connectomics reconstructions (Tavakoli et al., 2025), such local narrowing along spine necks is also apparent.

Our study illustrates how dense EM reconstructions can be transformed into functional, biophysically interpretable models. Rather than treating connectomics as purely structural, we used EM-derived morphologies and spine statistics to build data-driven cable and compartmental models that expose how nano-scale geometry of spines controls the timing of synaptic integration and the efficacy of inhibition. Moving from EM-meshes and synapse coordinates to explicit predictions about voltage, timing windows, and gating indices is a step toward a broader “biophysics of connectomics”, in which dense reconstructions are routinely translated into their digital twin at the single-cell and network-level models incorporating realistic connectivity, ion channels, synaptic dynamics, and plasticity rules.

Future work should embed EM-based spine morphologies in fully active single-neuron and network models to test how nanoscale timing control at individual spines shapes population coding and learning. Coupling these biophysical models to deep neural-network “digital twins” at both the single-neuron level (Aizenbud et al., 2025) and network level (Wang et al., 2025) will allow systematic exploration of how *R*_*neck*_-dependent temporal filtering and DiS-based inhibitory gating affect task performance and interference-resistant learning. In particular, such models can address how many local gates on DiSs are required to expand the repertoire of context-dependent functions at both single-neuron and network levels; how DiSs might selectively route information from specific dendritic subtrees to the soma, implementing a single-cell analogue of attention, and how the sub-millisecond temporal precision of DiS gating enables neurons to selectively encode or suppress specific temporal patterns. These and related questions can be studied within a biologically grounded, unified *“biophysics of connectomics”* framework suggested in this study.

Finally, our theoretical predictions are directly testable using emerging experimental approaches. Fast voltage and calcium imaging at the level of individual spines can probe how EPSP duration, spikelet generation, and NMDA/Na^+^ channel recruitment depend on local neck geometry, spine density, and DiS configuration. Time-resolved optogenetic or uncaging paradigms that systematically vary E-I timing could assay the predicted temporal dependence of synaptic interaction on *R*_*neck*_. Across animals and learning paradigms, connectomic datasets could be re-analyzed to track how *R*_*neck*_ distributions, DiS prevalence, and spine density reorganize to test the possibility that temporal processing and inhibitory gating are themselves plastic targets of experience-dependent structural change.

## Methods

### Ultrastructural spine dataset

The full dataset includes 4,223 dendritic spines from layer 6 pyramidal cells in the somatosensory cortex of a young adult mouse (Kasthuri et al., 2015). Spines located on apical dendritic branches extending into layer 5 were manually segmented at a resolution of 6×6×30 nm, and their morphologies were previously analyzed (Ofer et al., 2021).

### Dendritic spines head-neck separation

To automatically separate dendritic spines into head and neck composition, we employed a previously developed algorithm based on computer vision tools (Ofer et al., 2021). The algorithm exploits the characteristic geometry of dendritic spines, in which a thin, approximately cylindrical neck connects a larger, approximately spherical head. Specifically, the spine mesh is partitioned based on local cross-sectional thickness along its length, enabling reliable identification of the narrow neck region and the expanded head region. However, in approximately 50% of the spines, the algorithm identified only one segment or more complex structures, rather than a distinct head and neck. This was particularly the case for stubby spines, filopodia, and branched spines. We therefore took only the spines with a well-separated head and neck (n = 2,074) to our morphological analysis (Figures 2 and 3).

From the perspective of electrical resistance estimation, the exact border between the head and neck is not very important, since the short sections connecting the neck and head are usually thick and contribute little to the overall resistance of the spine neck.

### Estimation of EM-based spine neck resistance – the “section-by-section” method

The classical method for measuring the electrical resistance of the neck assumes the neck is a cylindrical cable. Accordingly, spine neck resistance is proportional to the neck length divided by the average cross-sectional area, usually calculated using the averaged radius as *πr*^2^ (Equation 1).

In our novel approach, we used the spine neck skeleton, represented by tens of 3D points along the center of the spine neck, divided into nanoscale slices. For each pair of consecutive points, we calculated the normal vector between them and determined the orthogonal plane to this normal. We then computed the area enclosed between this perpendicular plane and the mesh of the neck (yellow area in Figure 1B). Importantly, these sections do not correspond to the physical EM slices, but instead represent virtual cross-sections derived computationally from the reconstructed 3D mesh. Using the distance, *l*_*i*_, between consecutive skeleton points as the length of each section, and assuming specific resistivity, *R*_*i*_, the resistance of this section was calculated as:

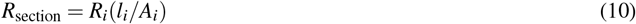

where *A*_*i*_ is the cross-sectional area of the section. By summing the (axial) resistances of all slices, modeled as series resistors, we obtained the total resistance of the spine neck (Equation 5).

To validate the method, we calculated the neck resistance for sections in both directions (from the dendritic shaft to the head and vice versa), which resulted in an average difference of only 4.3% in the total resistance across all modeled spines.

### Effective neck diameter calculation

The cross-sectional areas of sections along the spine neck often exhibit an elliptical shape (rather than a perfect circle). Therefore, we calculated an “effective diameter” based on the cross-sectional area, corresponding to the diameter of a circle with the same area, 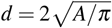. This method better reflects the electrical properties of the spine neck, as current flows through the neck and electrical resistance is primarily determined by the cross-sectional area.

### Quantifying the diameter irregularity along the spine neck

The normalized median absolute deviation (norm. MAD, Figures 2 and 3) was used to quantify the irregularity of diameters along the spine neck.

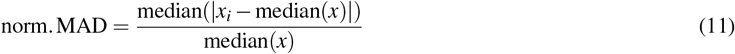

where *x*_*i*_ is the cross-sectional area of each nanoscale section along the spine neck. We used the median to mitigate the effect of extremely thin or thick slices and normalized the values to compensate for the total neck diameter in different spines.

### Equivalent cylinder for irregular dendritic spines

To avoid representing the individual spine by multiple serial compartments, we used an “equivalent cylindrical spine” model (Supplementary Figure S1). The equivalent cylinder of length *l* and radius *r* preserves both *R*_*neck*_ and the total surface area, *A*_*neck*_, of the EM-reconstructed dendritic spine, as computed by the section-by-section method (Equation 5). Namely,

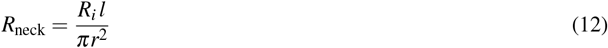

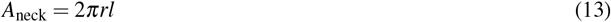

This yields closed-form expressions for the radius, *r*, and length, *l*, of the equivalent spine:

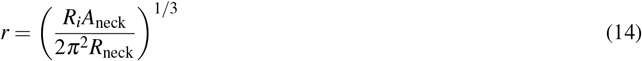

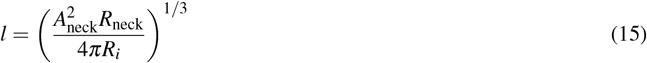

Supplementary Figure S1 shows that the voltage response at the spine head in the fully-modeled spine and in the equivalent spine model is very close. We therefore modeled all spines using the equivalent spine approximation.

### NEURON simulations

To investigate the functional implications of dendritic spine morphology, we performed compartmental simulations using NEURON (8.2.0) (Hines and Carnevale, 1997). All simulations were run using experimentally constrained passive membrane properties (*R*_*i*_ = 100Ω*cm, E*_*pas*_ = *™*68*mV, R*_*m*_ = 10, 000Ω*cm*^2^; *C*_*m*_ = 1*µF/cm*^2^). Unless stated otherwise, dendrites were modeled as an electrotonically long, uniform cylindrical cable. The modeled spine was located at the middle of the dendritic cable. Spatial discretization was set to Δ*X* = 0.02*λ*, and the integration time step was set to Δ*t* = 0.00025*ms*. Simulation duration was 30*ms* for single-input and 65*ms* for multi-input.

### Frequency and phase response at the spine head membrane

In Figures 3 and 5, frequency-domain analysis was computed using NEURON’s built-in ‘Impedance’ function. A small-amplitude sinusoidal current was injected at the spine head, and the resulting voltage was recorded there. For each input frequency, *f*, we computed the impedance (*Z*(*f*) = *V* (*f*)*/I*(*f*)) and phase, impedance magnitude was converted to dB relative to the lowest-frequency (*Z*(*f*_*min*_ = 0)):

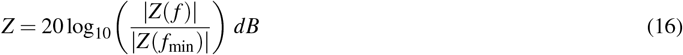

Cutoff-frequency is defined as *Z*(*f*_*c*_) = −3*dB*

### Synaptic input and receptor dynamics

Across all simulations, synaptic inputs were implemented using the following models. For *AMPA*-based excitatory and *GABA*_*A*_-based inhibitory synapses, the synaptic current, *I*_*syn*_(*t*), and conductance *g*_*syn*_(*t*) are,

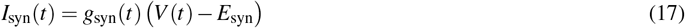

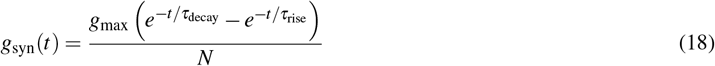

where *g*_*max*_ is the maximal synaptic conductance, *τ*_*decay*_ and *τ*_*rise*_ are the decay and rise time constants, respectively, and *N* is a normalization constant,

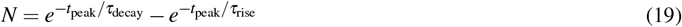

where *t*_*peak*_ is the peak-time of the synaptic conductance. For NMDA:

For NMDA:

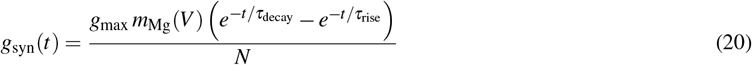

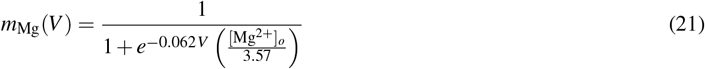

In all figures, we used for the AMPA-based excitatory synapse with *τ*_*rise*_ = 0.1*ms, τ*_*decay*_ = 1*ms. g*_*max*_ = 0.4*nS*, and *E*_*AMPA*_ = 0*mV* in Figures 3 and 5, and *g*_*max*_ = 1*nS* in Figures 6 and 7. In Figures 6 - 8, the *GABA*_*A*_-like synapse was modeled with *τ*_*rise*_ = 0.5*ms, τ*_*decay*_ = 5*ms, g*_*max*_ = 3.3*nS*, with *E*_*GABA*_ = *™*68*mV* (equal to the resting membrane potential). Supplementary simulations (Figures S2-S7) included two additional variants. First, to test the effect of fast inhibition, we repeated the simulations using the above AMPA-based kinetics also for the inhibitory conductance; namely, with *τ*_*rise*_ = 0.1*ms, τ*_*decay*_ = 1*ms* (all other parameters unchanged). In Figure 8, we used an AMPA- and NMDA-based excitatory synapse: for AMPA: *τ*_*rise*_ = 0.5*ms, τ*_*decay*_ = 3*ms, g*_*max*_ = 1*nS*, and *E*_*AMPA*_ = 0*mV*. For NMDA: *τ*_*rise*_ = 3*ms, τ*_*decay*_ = 30*ms, g*_*max*_ = 2*nS*.

For the simulations where we kept the peak EPSP identical for all *R*_*neck*_ cases (Supplementary Figures S3, S4, and S7), we adjusted the excitatory AMPA peak conductance 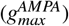 separately for each *R*_*neck*_, such that the EPSP peak at the spine head reached *™*35*mV*. This tuning was performed via a binary search algorithm (with 0.1*mV* tolerance). In the corresponding inhibition simulations (E+I case), inhibitory strength was set proportionally as: 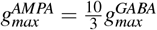.

NMDA-mediated calcium current was estimated with Goldman-Hodgkin-Katz (GHK) current equations:

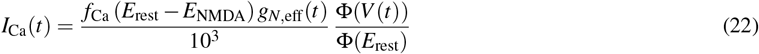

The calcium component of the NMDA current was approximated as a fixed fraction of the total NMDA current, *f*_*Ca*_ = 0.1, consistent with experimental measurements of *Ca*^2+^ permeability of NMDA receptors (Jahr and Stevens, 1993). Here, *E*_*rest*_ = −68*mV, E*_*NMDA*_ = 0*mV*, and *g*_*NMDA,eff*_ in nS.

The voltage-dependent GHK flux factor was computed as:

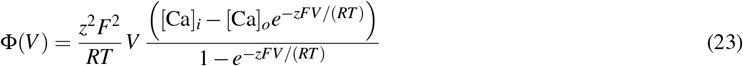

where *Ca*_*i*_ and *Ca*_*o*_ denote the intracellular and extracellular *Ca*^2+^ concentrations, respectively; *z* = 2 is the valence of *Ca*^2+^; *F* is Faraday’s constant; *R* is the universal gas constant; and *T* is the absolute temperature.

Free calcium concentration was estimated using:

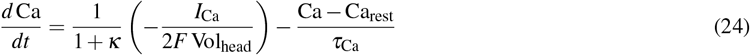

We set the spine head volume, *Vol*_*head*_, to 0.027*µm*^3^; 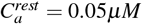 and *τ*_*Ca*_ = 20*ms. F* is Faraday’s constant, and *κ* = 400 is an effective buffering factor.

### F-Factor approximation for incorporating dendritic spines into the dendritic cable

In Figure 5, we varied the spine density along the modeled dendrite to capture the aggregate effect of the impedance load on the temporal dynamics at individual spines. To avoid modeling all spines for each density case, we incorporated all spines (except the one receiving the input) into the dendritic cable using the F-factor approximation (Holmes and Woody, 1989; Rapp et al., 1992) whereby:

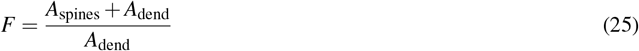

where *A*_*dend*_ is the membrane area of the dendritic cable, and *A*_*spines*_ is the total membrane area of all spines. The F-factor is then used to scale the specific cable parameters such that 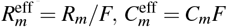. We validated this approximation by comparing responses from explicit multi-spine models to the respective approximation using the F-factor, confirming matching of impedance and voltage waveforms at the modeled spine for the two cases.

## Code availability

The code used in this study is available at *GitHub*: https://github.com/NetanelOfer/Spine_timing. The 3D meshes of the analyzed dendritic spines are available at https://doi.org/10.7916/d8-tdqd-dh88.

## Acknowledgements

We thank all members of Segev’s lab and members of the NIH “Dendritic Consortium” — Jayeeta Basu, Michael Lin, Jeff Lichtman, Elly Nedivi, and Rafael Yuste — for fruitful discussions and valuable feedback regarding this work. The work was supported by the NINDS 1RM1NS132981-01 grant.

## Appendix A Section-by-section estimation yields higher *R*_neck_ than averaging

In the uniform-cylinder approximation, the spine-neck resistance (*R*_neck_), with neck length ℓ, radius *r*, and specific axial resistivity *R*_*i*_ (in Ω·cm) is:

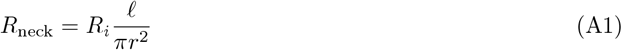

In practice, real spines do not have a uniform neck diameter. As a first departure from the uniform-cylinder model, we consider a spine neck composed of two cylindrical segments with radii *a < b* and lengths ℓ_*a*_ and ℓ_*b*_, respectively (Figure A1), and later extend the analysis to necks with more than two radii (multiple segments).

**Figure A1.**
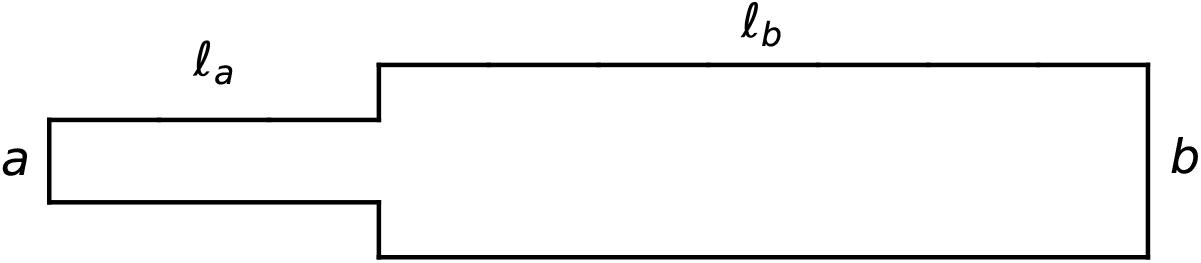
Spine neck composed of a small-diameter segment *a* of length ℓ_*a*_ and a large-diameter segment *b* of length ℓ_*b*_.

Traditionally, *R*_neck_ is estimated using the “average method”, which relies on the total length (ℓ_*a*_ + ℓ_*b*_) of the spine and a weighted average of the radii *a* and *b* (Figure A2, left).

**Figure A2.**
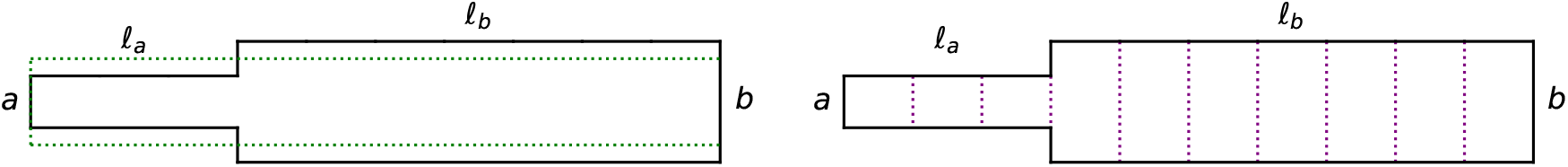
Left: The “average” method to estimate *R*_neck_ (weighted average is depicted in green; Equation A2). Right: The “section-by-section” method to estimate *R*_neck_ (sections are depicted in purple; Equation A3).

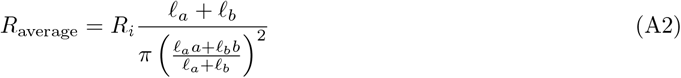

In contrast, our “section-by-section method” estimates the resistance by discretizing the neck into thin slices and summing their individual contributions along the entire neck length (Figure A2, right).

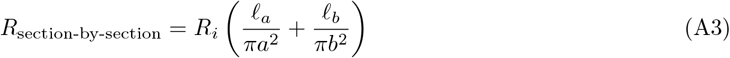

where *R*_*i*_ (in units of Ω*cm*) is the specific axial resistivity.

We illustrate hereby the derivation of *R*_neck_ for only two regions along the spine neck (radii *a, b* and lengths ℓ_*a*_, ℓ_*b*_), the same argument applies to any partition of the neck into *n* regions with radii *r*_*i*_ and lengths ℓ_*i*_.

### Proof of the Inequality - *R*_average_ ≤ *R*_section-by-section_

To prove that *R*_average_ ≤ *R*_section-by-section_ for all positive ℓ_*a*_, ℓ_*b*_, *a*, and *b*. It suffices to show that:

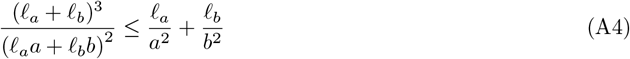

where ℓ_*a*_, ℓ_*b*_, *a*, and *b* are positive real numbers.

#### Step 1: Define Weight Variables

Let us define weight variables *m*_1_ and *m*_2_ as:

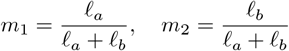

These weights satisfy:

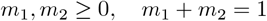

#### Step 2: Rewrite the Inequality Using Weights

Express the inequality in Equation A4 in terms of *m*_1_ and *m*_2_:

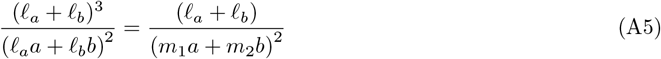

Similarly, the right-hand side becomes:

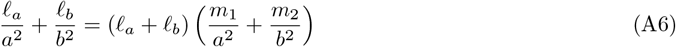

Thus, the inequality simplifies to:

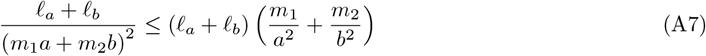

Dividing both sides by ℓ_*a*_ + ℓ_*b*_:

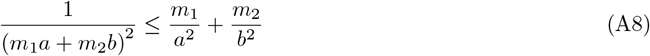

#### Step 3: Jensen’s Inequality

Jensen’s inequality is a fundamental result in convex analysis. It states that for a convex function *ϕ*(*x*) and a set of weights *m*_1_, *m*_2_, …, *m*_*n*_ such that *m*_*i*_ ≥ 0 and 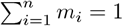, we have:

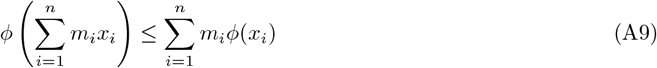

#### Step 4: Apply Jensen’s Inequality

With 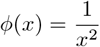 convex on *x >* 0 (since 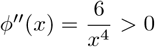), Jensen gives *ϕ* (*m*_1_ *a* + *m*_2_ *b*) ≤ *m*_1_ *ϕ*(*a*) + *m*_2_ *ϕ*(*b*) (here n=2).

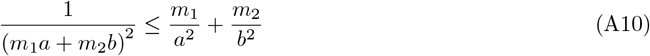

#### Step 5: Conclude the Proof

Multiplying both sides by ℓ_*a*_ + ℓ_*b*_:

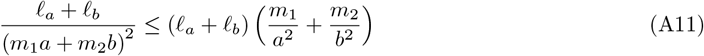

Recall that:

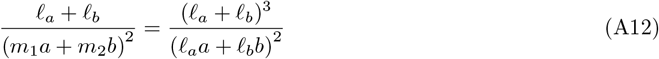

and

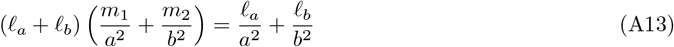

Therefore:

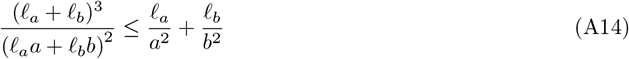

#### General Case: A Cable with Multiple Diameters

Let 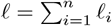 Then

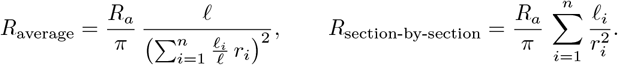

Since *ϕ*(*x*) = 1*/x*^2^ is convex for *x >* 0, applying Jensen’s inequality with weights ℓ_*i*_*/*ℓ and points *r*_*i*_ gives *R*_average_ ≤ *R*_section-by-section_ for any *n*.

## Appendix B The Effect of Spine Neck Resistance on System Time Constants

### 1 Introduction

A point neuron can be modeled as a single RC circuit – a resistor and capacitor in parallel – where a single membrane time constant, *τ*_*m*_ = *RC*, determines the temporal evolution of the membrane voltage *V* (*t*) in response to input currents (Lapicque, 1907; Hodgkin and Huxley, 1952; Rall, 1957).

In contrast, dendrites behave as distributed electrical cables that can be approximated as a chain of coupled RC elements, such that *V* (*t*) is governed by multiple time constants (*τ*_*m*_, *τ*_1_, *τ*_2_, …). These additional ‘equalizing’ time constants (*τ*_1_, *τ*_2_, …) accelerate voltage dynamics, yielding an effective time constant that is shorter than that of an equivalent lumped RC compartment (Rall, 1957, 1960, 1969).

Here, we show that a dendritic spine – whose head membrane is modeled as an RC circuit – connected to its parent dendrite through a variable resistor representing the spine neck (whose value depends on neck morphology) can exhibit an ultra-fast effective time constant at the spine head (Figure B1A).

**Figure B1.**
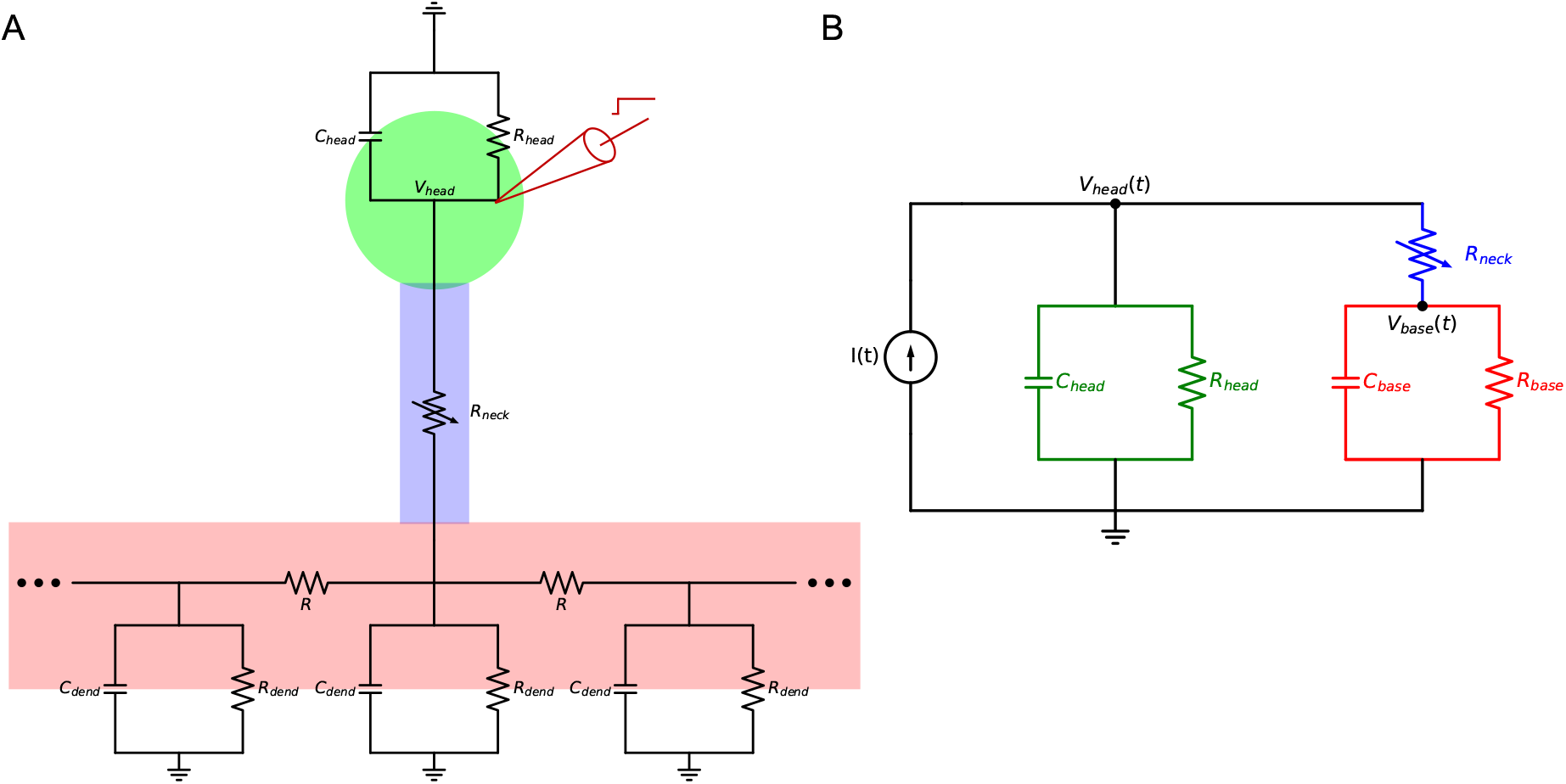
Electrical circuit representation of a dendritic spine. **A**. Circuit schematics: spine head (green), spine neck (blue), and dendritic shaft (red). **B**. Lumped-circuit diagram of the circuit in A, where the dendrite is modeled as a single RC circuit coupled to the spine base. A step current is applied to the spine head (red electrode)

First, we analytically solved the lumped-circuit model (Figure B1B) to derive expressions for the two time constants and their respective coefficients that govern *V* (*t*) (Section 2). We then performed numerical simulations in SPICE on the distributed model (Figure B1A) to obtain *V* (*t*) at the spine head as a function of *R*_neck_, thereby capturing the combined effect of all time constants on the voltage dynamics at the spine head membrane for the full circuit (Section 3).

### 2 Lumped-Circuit Analysis

The lumped circuit consists of a step current injection *I*(*t*) at the spine head, modeled by an RC element (*R*_head_, *C*_head_), in parallel with a branch comprising the spine-neck resistance *R*_neck_ and the dendritic shaft RC element (*R*_base_, *C*_base_). We assume identical membrane properties at the spine head and the spine base, such that *R*_head_*C*_head_ = *R*_base_*C*_base_ = *τ*_*m*_.

Applying Kirchhoff’s Current Law (KCL) gives the following governing equations:

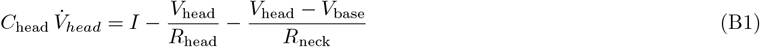

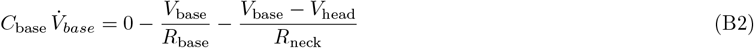

In matrix form 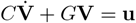 where,

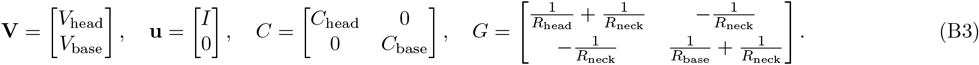

At *t* → ∞, capacitors behave as open circuits 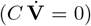, so the steady-state voltages satisfy *G* **V**_∞_ = **u**. The solution is given by **V**_∞_ = *G*^−1^**u**, where

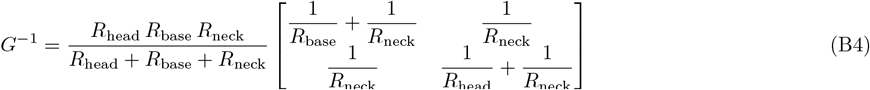

Multiplying *G*^−1^ by the input vector (*I*, 0)^T^ yields the steady-state voltage response at the spine head and base:

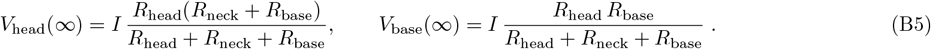

It is intuitive to see that the injected current *I* splits between two parallel branches: the “head” branch (*R*_head_) and the combined “neck + base” branch (*R*_neck_ + *R*_base_). Accordingly, the voltage at the spine head, *V*_head_, is given by the product of the total current and the equivalent parallel resistance, *R*_head_ ∥ (*R*_neck_ + *R*_base_). Within the “neck + base” branch, the voltage at the spine base, *V*_base_, is obtained by applying the voltage divider between *R*_neck_ and *R*_base_ to the head potential *V*_head_.

We next normalize all subsequent voltages to the steady-state value *V*_head_(∞).

**Homogeneous dynamics and time constants.** Let **v**(*t*) = **V**(*t*) *™***V**_∞_. Then 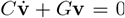. Substituting an exponential solution **v**(*t*) = **v**_0_*e*^*λt*^ yields the eigenvalue condition (*λC* + *G*)**v**_0_ = 0, so the decay rates are obtained by setting the determinant to zero:

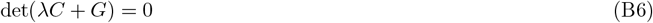

which is a quadratic in *λ*. Hence there are two negative real roots, *λ*_1_ and *λ*_2_, corresponding to the two exponential modes. Under the matched RC assumption, one eigenvalue can be obtained directly by testing the common–mode vector [1, 1], which represents both voltages varying equally (no current through *R*_neck_):

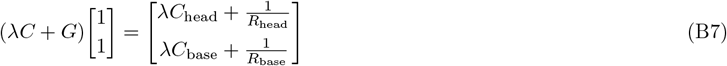

Both entries vanish simultaneously when

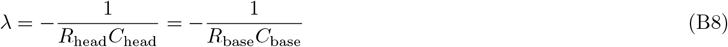

With *R*_head_*C*_head_ = *R*_base_*C*_base_ = *τ*_*m*_, this gives

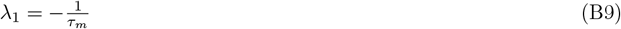

Corresponding to the time constant:

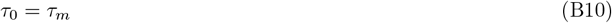

Multiplying 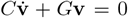 by *C*^−1^ gives 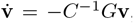, so the decay rates are the eigenvalues of −*C*^−1^*G*. Alternatively, the eigenvalues can be obtained from the trace and determinant of *™C*^−1^*G*. Once *λ*_1_ is known, the second eigenvalue *λ*_2_ follows directly from the trace identity:

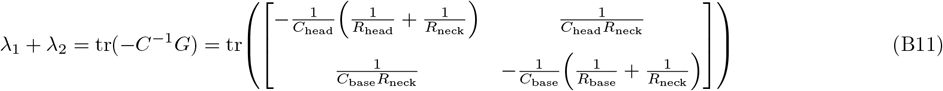

Evaluating the trace explicitly yields:

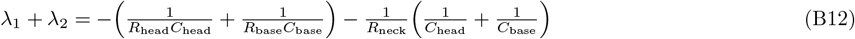

Inserting *λ*_1_ = −1*/τ*_*m*_ and *R*_head_*C*_head_ = *R*_base_*C*_base_ = *τ*_*m*_ gives

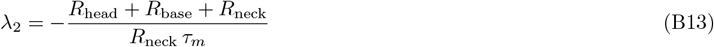

Therefore the second time constant is

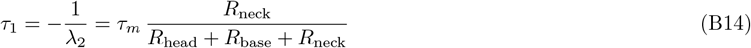

Equation B14 shows that *τ*_1_ is smaller than *τ*_*m*_, depends on *R*_*neck*_, and converges to *τ*_*m*_ as *R*_*neck*_ becomes large. Consequently, the normalized head voltage can be expressed as a sum of two *single–pole step responses*:

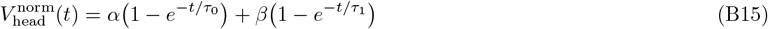

The coefficients *α* and *β* are determined from the normalized dynamics as follows.

(1) Because 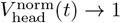 as *t* →*∞*, it follows that *α* + *β* = 1.

(2) The initial normalized slope refers to the time derivative of 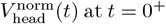 at *t* = 0^+^, after normalizing the voltage to its steady-state value. Differentiating (B15) and evaluating at *t* = 0 gives:

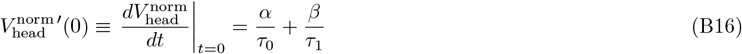

From (B1), at *t* = 0^+^ the capacitors are uncharged, so 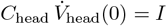, where *V*_∞_ is taken from (B5), and hence:

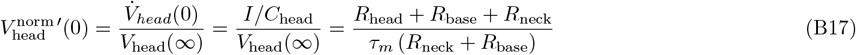

By equating the two expressions for the initial normalized slope, (B16) and (B17), we obtain:

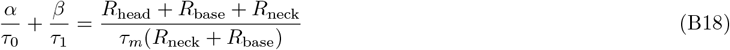

Substituting *τ*_0_ from (B10) and *τ*_1_ from (B14), we obtain

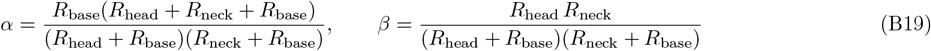

Inserting the values of *α* and *β* from (B19) into (B15), we obtain the explicit time-domain solution:

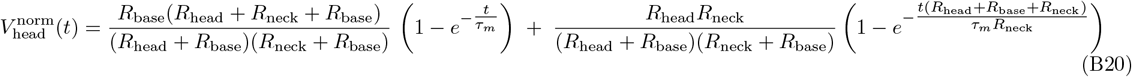

#### 2.1 Biophysically-Realistic Parameters Predicts Sub-Millisecond Effective Time Constants

We assume identical membrane properties at the spine head and the spine base, such that *R*_head_*C*_head_ = *R*_base_*C*_base_. The small membrane area of the spine head yields high resistance and low capacitance, whereas the much larger spine base yields small *R*_*base*_ and large *C*_*base*_. We set the membrane time constant for both compartments to *τ*_*m*_ = 10 ms, consistent with experimentally measured values (Koch et al., 1996), using the following parameter values:

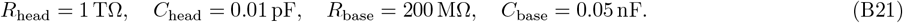

We used these values in Equations (B10) and (B14) for various values of *R*_neck_. *τ*_0_ is independent of *R*_neck_ and remains fixed at *τ*_*m*_ = 10 ms. In contrast, *τ*_1_ is very small at low *R*_neck_ and increases with *R*_neck_ (Figure B2A). To assess the relative contributions of *τ*_0_ and *τ*_1_ to the effective time constant, we computed *α* as a function of *R*_neck_ (Equation B19; Figure B2B). The effective time constant, *τ*_eff_, is defined as the time at which the spine-head voltage response (Equation B20) reaches 63% of its steady-state value (Figure B2C). Note that *τ*_eff_ decreases steeply already for relatively small *R*_neck_ values, reaching a minimum of ≈ 200 *µ*s at *R*_neck_ around 1 GΩ. This reduction in *τ*_eff_ gives rise to ultra-fast (sub ms) voltage response at the spine head membrane.

**Figure B2.**
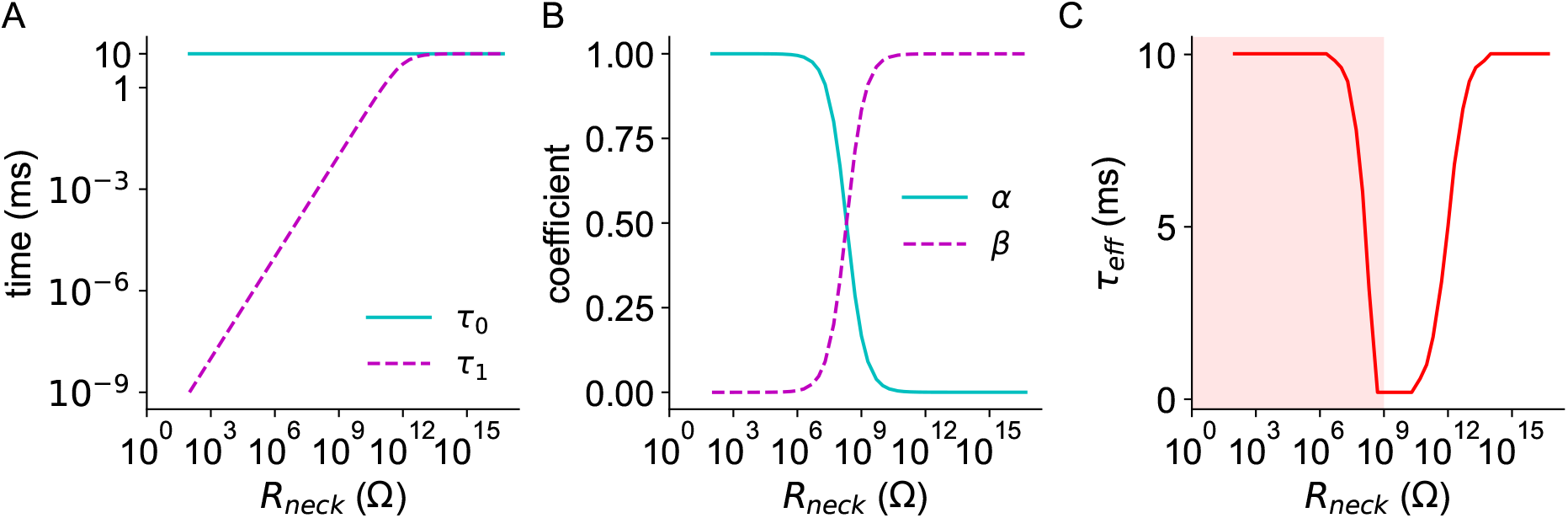
The effect of spine neck resistance on the temporal dynamics of the spine head membrane. **A**. The two time constants, *τ*_0_ and *τ*_1_, as a function of *R*_neck_, calculated by substituting physiological parameter values into Equations (B10) and (B14). **B**. The coefficients *α* (and *β* = 1 − *α*) that determine the relative contributions of the two exponential components governed by *τ*_0_ and *τ*_1_, respectively, as a function of *R*_neck_ (Equation B19). **C**. Effective time constant, *τ*_eff_, defined as the time for *V*_head_ to reach 63% of its steady-state value in response to a step current input, as a function of *R*_neck_.

Although the analysis extends to *R*_neck_ values far above those found in biological systems, its purpose is to illustrate the phenomenon in general terms. Notably, the biologically relevant range (0–1GΩ) (Harnett et al., 2012; Cornejo et al., 2022) remains within the region where the chosen parameters produce a substantial decrease in delay time (shaded area in Figure B2C).

### 3 Numerical Simulations of the Full Circuit

To evaluate the effective time constant (*τ*_eff_) of the full circuit (Figure B1A) as a function of *R*_neck_, we performed SPICE simulations to analyze the voltage response in the spine head to a step current injection. In these simulations, the dendrite was represented by 1,000 segments (each modeled as an RC circuit), such that:

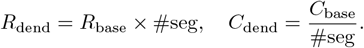

where *R*_base_ and *C*_base_ are as shown in B21.

The two extreme cases, *R*_neck_ → ∞ and *R*_neck_ = 0, were previously analyzed by Rall (1957, 1960). In the limiting case of *R*_neck_ → ∞, the spine he_(_ad becom_)_es effectively isolated, and its dynamics are those of an isopotential soma, with the voltage rising as *V*_*norm*_(*t*) =1 − *e*^−*t/τ*^ (gray dash-dotted curve in Figure B3A). Conversely, when *R*_neck_ = 0, the current is injected directly to the dendritic shaft, and the voltage follows 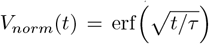 (black dashed curve in Figure B3A). Our results further reveal that the voltage dynamics for intermediate values of *R*_neck_ relevant to dendritic spines, specifically *R*_neck_ = 100 MΩ and *R*_neck_ = 1 GΩ (Figure B3A), exhibit a substantially more rapid voltage response compared to both limiting cases. Figure B3B shows the effective time constant, *τ*_*eff*_, normalized by the membrane time constant, *τ*_*m*_, as a function of *R*_neck_, revealing a distinctive biphasic (U-shaped) behavior, in which *τ*_*eff*_ first decreases and then increases as neck resistance rises.

**Figure B3.**
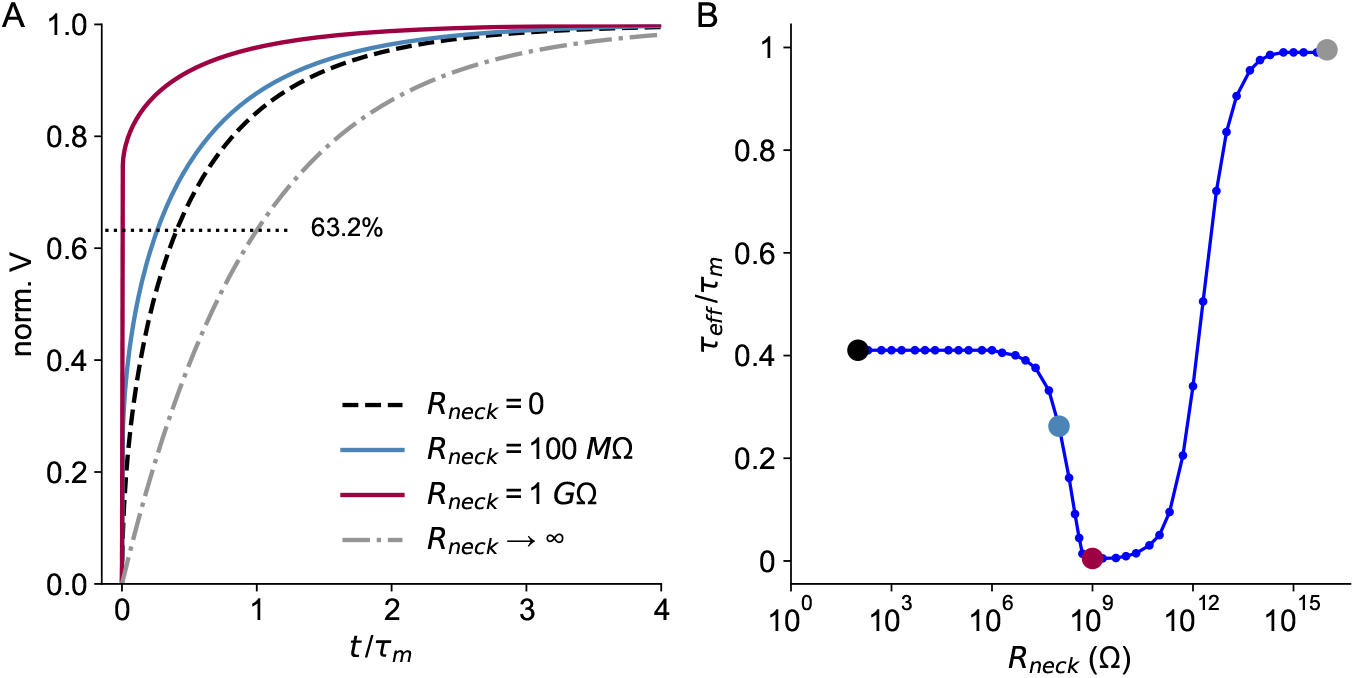
Effective time constant in a spine coupled to a dendritic cable as a function of *R*_neck_. **A**. Example voltage traces for several *R*_neck_ values in response to a step current, showing the effect of neck resistance on the voltage dynamics. **B**. Normalized effective time constant (*τ*_eff_) as a function of *R*_neck_, defined as the time required to reach 63% of the steady-state voltage. The colors of the marked data points correspond to the voltage traces shown in panel A.

### 4 Biophysical Interpretation of the Biphasic Dependence of *τ*_*eff*_

To better understand why the effective membrane time constant first shortens and then lengthens as *R*_neck_ increases, we plotted the relevant components in Figure B4. The first contributing factor is the relative weighting between the two exponential components that determines the voltage response (see Equation B20). This weighting, denoted by *α* (Equation B19; dashed green curve in Figure B4), determines which of the two system time constants (*τ*_0_ and *τ*_1_) dominates *V* (*t*). For small values of *R*_neck_, the voltage dynamics are governed primarily by the slower component. As *R*_neck_ increases, the faster component becomes increasingly dominant, which causes the effective time constant to decrease around *R*_neck_ ≈ 10^8^Ω.

**Figure B4.**
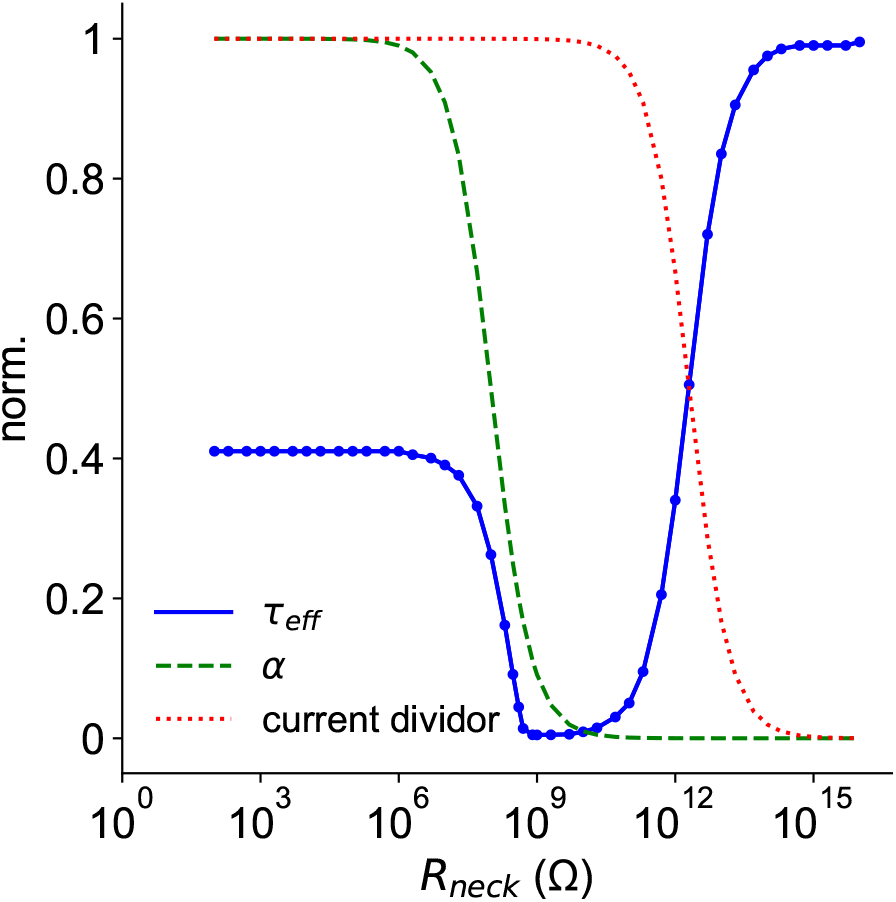
**Dependence of the effective time constant on** *α* **and current division as a function of** *R*_neck_ for the circuit shown in Figure B1A. The dashed green curve shows the coefficient, *α* - the relative weight of the fast (*τ*_1_) and slow (*τ*_0_) time constants that govern the voltage response in this system. The dotted red curve shows the fraction of apllied current flowing through the spine neck. Parameter values are the same as in Subsection 2.1 (Equation B21). Together, these curves illustrate how the two mechanisms jointly determine the biphasic dependence of the effective membrane time constant (blue) on *R*_neck_.

The second factor arises from the current division between the “head” branch and the combined “neck + base” branch (see Figure B1B). The fraction of current flowing through the neck is described by the current divider (Equation B22; dotted red curve in Figure B4). When *R*_neck_ is small, most of the current escapes through the “neck + base” pathway. At very high *R*_neck_ (around 10^12^Ω), the current becomes largely restricted to the spine head compartment, and the effective time constant approaches the intrinsic membrane time constant of the head, becoming essentially independent of the weighting factor *α*. Together, these two mechanisms, acting over different ranges of *R*_neck_, give rise to the biphasic (U-shaped) dependence of the effective time constant on *R*_neck_.

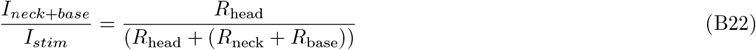

### 5 Frequency-Domain (AC) Analysis of the Lumped Equivalent Circuit

To analyze the voltage response to varying current input frequencies, we performed an AC analysis for the lumped circuit (Figure B1B).

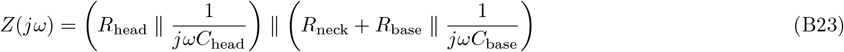

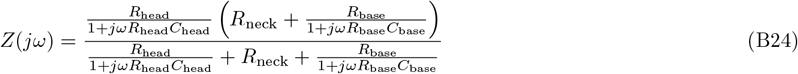

Numerical substitution with parameter values (same as in Subsection 2.1; Equation B21) gives:

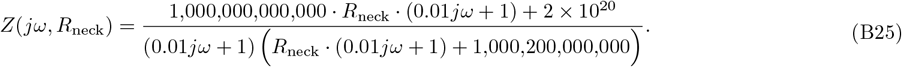

From *Z*(*jω*), we generated the Bode plot (Figure B5A,B), showing magnitude and phase as functions of frequency for three representative values of *R*_neck_ (0, 100 MΩ, and 100 TΩ).

**Figure B5.**
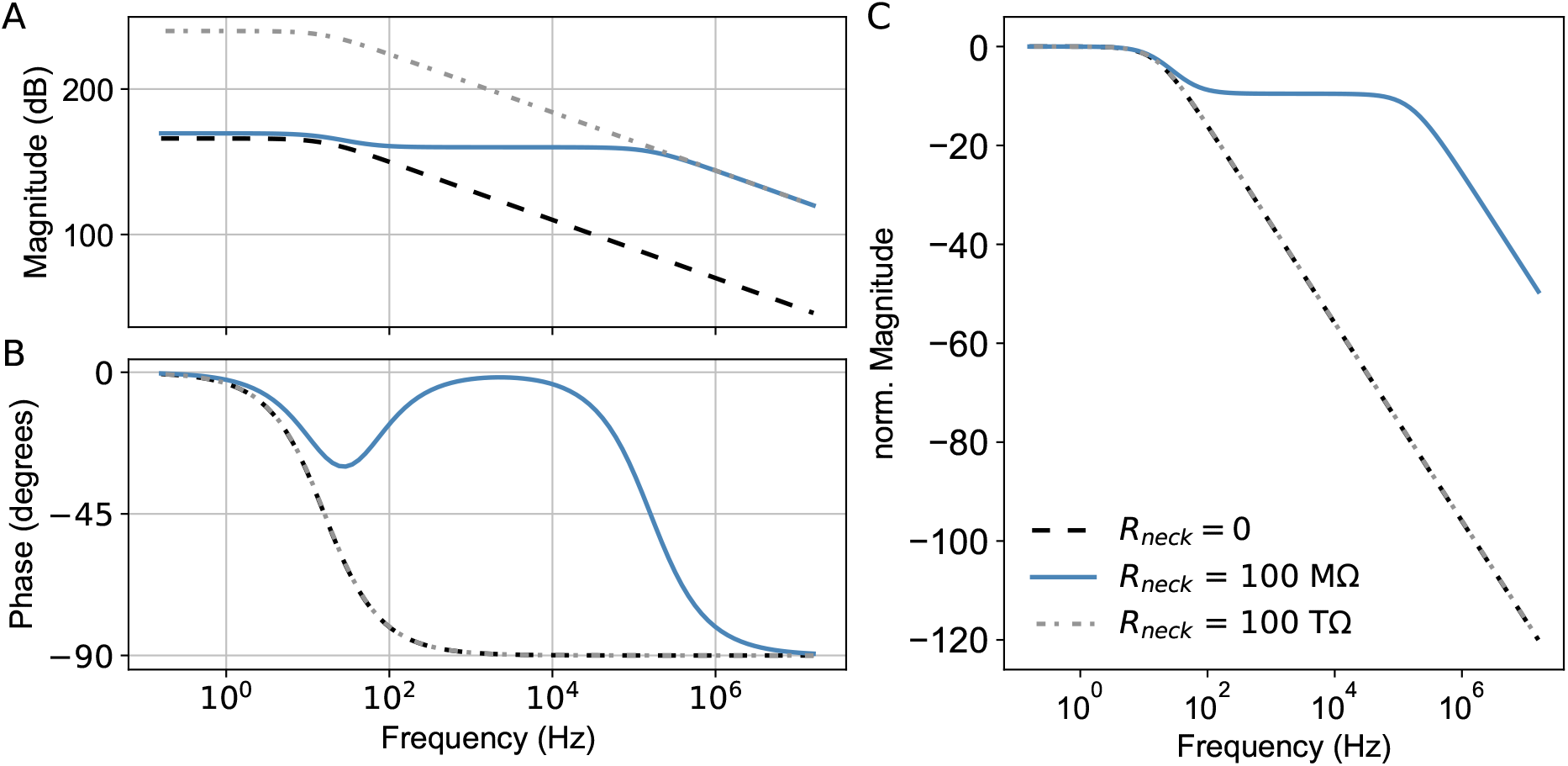
**A-B**. Bode plot showing the magnitude and phase responses for different values of *R*_neck_, derived from Equation B25. **C**. Normalized magnitude response (in dB) relative to the DC value. The two extreme cases (*R*_neck_ = 0 and *R*_neck_ →∞) show nearly identical profiles due to normalization. In contrast, for dendritic spines with intermediate neck resistances (e.g., 100 MΩ), the response remains elevated at higher frequencies.

We examined how varying *R*_neck_ affects the response across a range of frequencies. Extremely low and high *R*_neck_ values correspond, respectively, to current applied directly to the dendritic shaft or to spines with very high neck resistance. Our results show that dendritic spines can preserve a high response magnitude at higher frequencies without substantially altering the phase (Figure B5C).

### 6 Code and Simulation Availability

All analytic code and SPICE simulation files (implemented in Python with the PySpice package, v1.5) used to generate the results are publicly available at: https://github.com/NetanelOfer/Spine_timing

---

**Figure S1.**
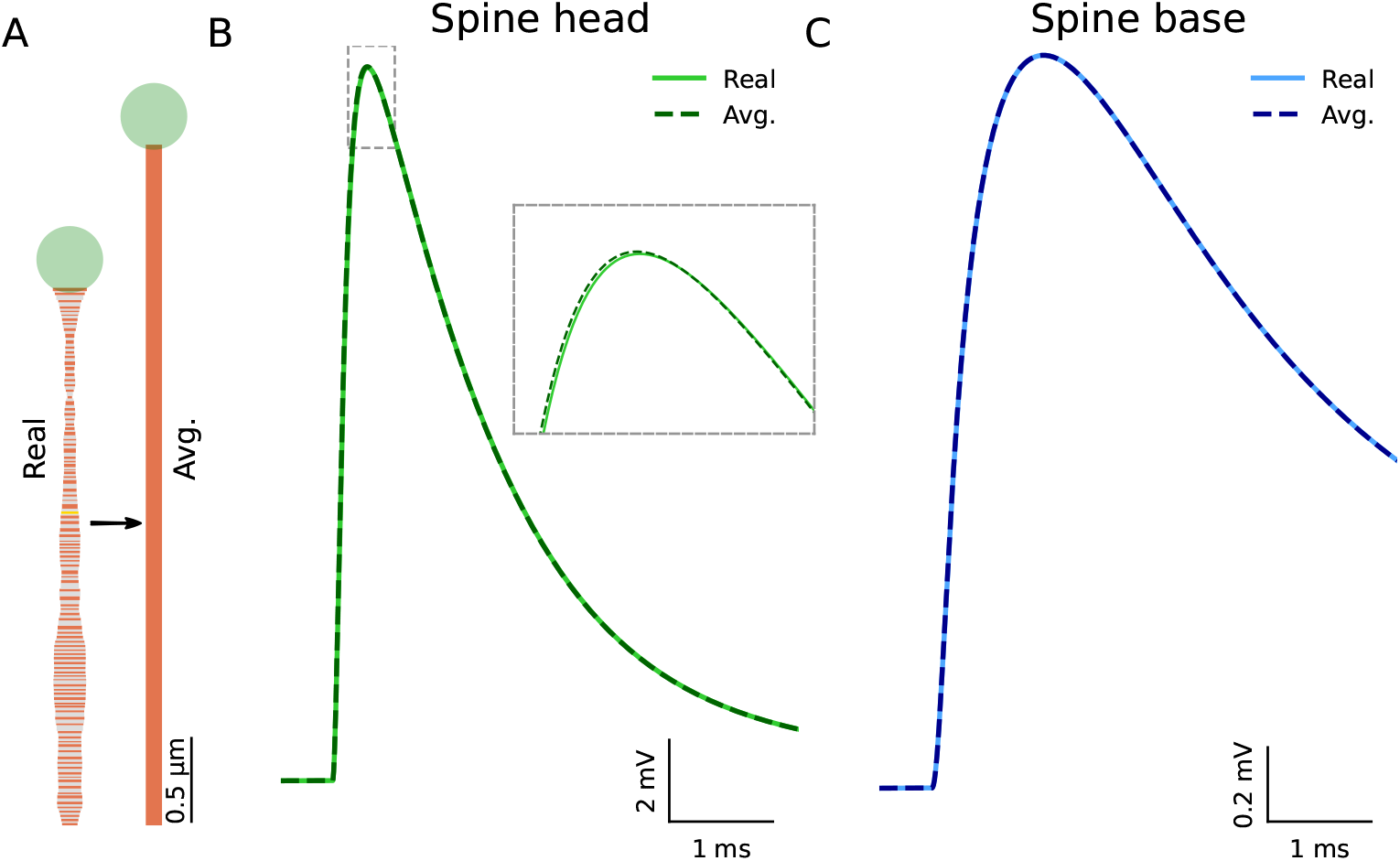
An equivalent cylindrical cable to an irregular spine. **A**. Left: Section-by-section representation of an EM-reconstructed spine with *R*_*neck*_ = 800*M* Ω (as in Figure 1C). Right: The respective “equivalent cylindrical spine”. **B-C**. Comparison of the EPSP at the spine head (B) and base (C) in the section-by-section versus the equivalent cylinder computation. Model consists of a 1*µm* diameter 25*λ* long passive cable (1005 segments) with *C*_*m*_ = 1*µF/cm*^2^; *R*_*m*_ = 10, 000Ω*cm*^2^ (resulting in *τ*_*m*_ = 10*ms*) and *R*_*i*_ = 150Ω*cm*. Synaptic parameters: *E*_*AMPA*_ = 0*mV*; *g*_*max*_ = 1*nS* with *τ*_*rise*_ = 0.1*ms* and *τ*_*decay*_ = 1*ms*. Equivalent spine neck radius is 0.0486*µm*, and the length is 4.08*µm*.

**Figure S2.**
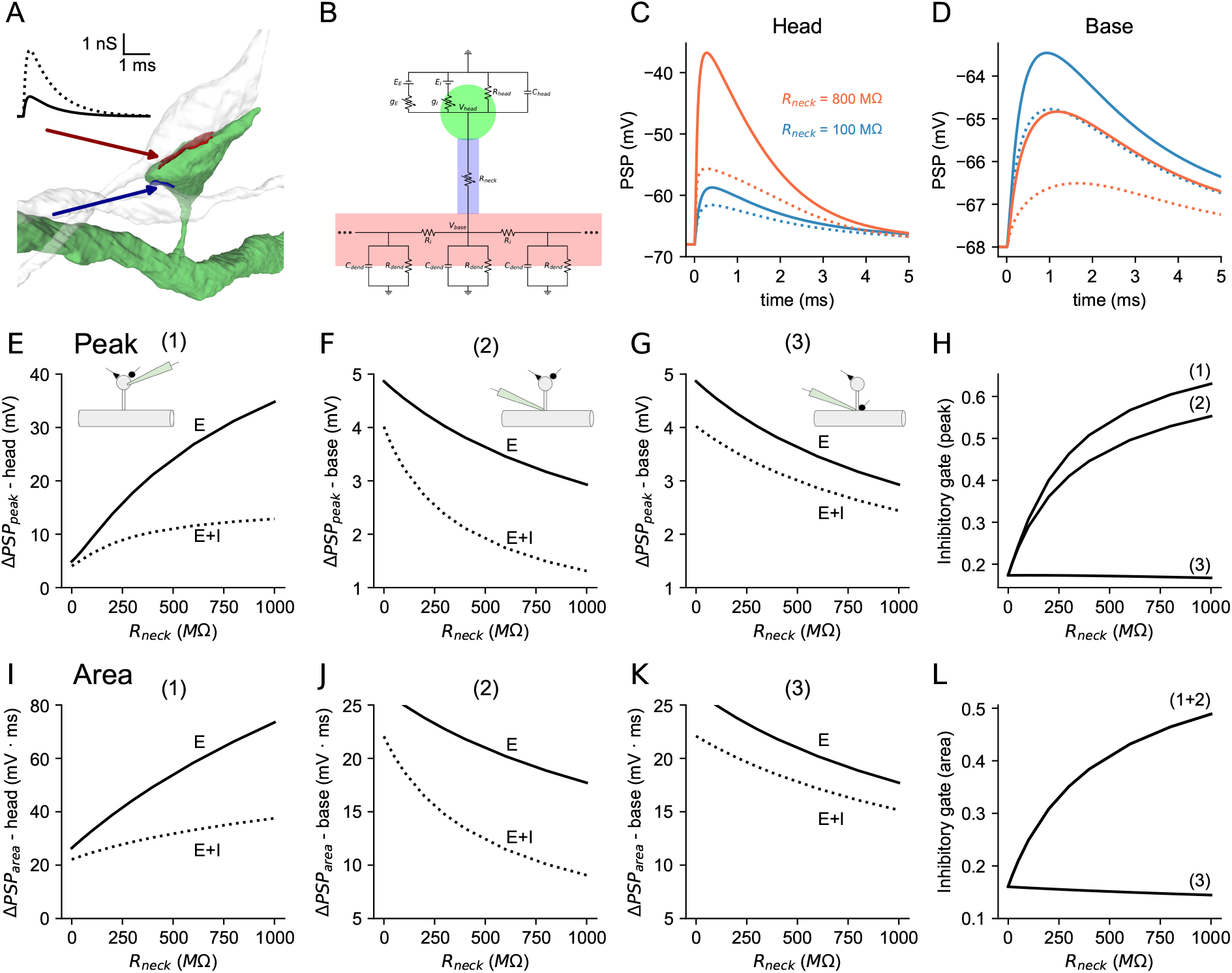
Potent inhibitory gating in dually innervated dendritic spines (DiSs) with AMPA-like inhibition. **A**. An EM-reconstruction of DiS. The two synapses are shown in red and blue (and with respective arrows). The inset shows the amplitude and time-course of the excitatory (continuous line) and inhibitory (dotted line) conductances. **B**. Equivalent circuit for a DiS on a near-infinite passive dendritic cable. **C**. PSPs at the spine head with (dotted line) and without inhibition (continuous line) for *R*_*neck*_ = 100*M* Ω (blue) and 800*M* Ω (orange). **D**. As in C, but the resultant PSPs at the spine base are shown. **E**. PSP peak at the spine head without (E) and with (E + I) inhibition as a function of *R*_*neck*_. **F**. As in E, but at the spine base. **G**. PSP peaks at the spine base when E is at the spine head and I at the spine base. **H**. Inhibitory gate (Equation 7) for the cases shown in E-G. **I-L**. Similar to E-H, but for area. Model parameters: AMPA: *τ*_*rise*_ = 0.1*ms, τ*_*decay*_ = 1*ms, g*_*max*_ = 1*nS*. GABA: *τ*_*rise*_ = 0.1*ms, τ*_*decay*_ = 1*ms, g*_*max*_ = 3.3*nS. E*_*GABA*_ = −68*mV* (equals the resting potential).

**Figure S3.**
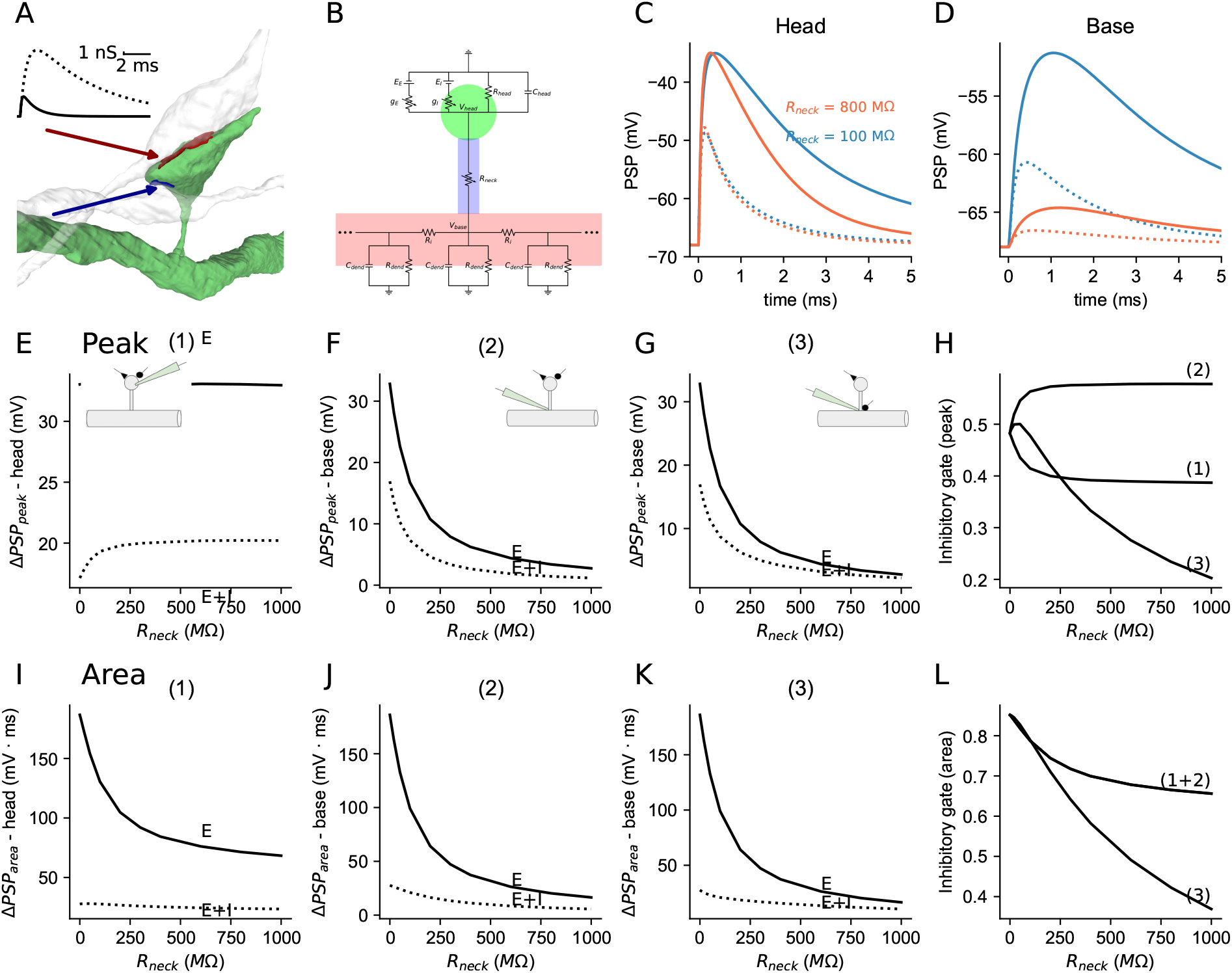
Potent inhibitory gating in dually innervated dendritic spines (DiSs) with −35 mV EPSP peak. **A**. An EM-reconstruction of DiS. The two synapses are shown in red and blue (and with respective arrows). The inset shows the amplitude and time-course of the excitatory (continuous line) and inhibitory (dotted line) conductances. **B**. Equivalent circuit for a DiS on a near-infinite passive dendritic cable. **C**. PSPs at the spine head with (dotted line) and without inhibition (continuous line) for *R*_*neck*_ = 100*M* Ω (blue) and 800*M* Ω (orange). **D**. As in C, but the resultant PSPs at the spine base are shown. **E**. PSP peak at the spine head without (E) and with (E + I) inhibition as a function of *R*_*neck*_. **F**. As in E, but at the spine base. **G**. PSP peaks at the spine base when E is at the spine head and I at the spine base. **H**. Inhibitory gate (Equation 7) for the cases shown in E-G. **I-L**. Similar to E-H, but for area. Model parameters: AMPA: *τ*_*rise*_ = 0.1*ms, τ*_*decay*_ = 1*ms*. GABA: *τ*_*rise*_ = 0.5*ms, τ*_*decay*_ = 5*ms, E*_*GABA*_ = −68*mV* (equals the resting potential). *g*_*max*_ = 1*nS* for AMPA and GABA were automatically calculated such that peak EPSP (AMPA only) will be equal to −35 mV peak.

**Figure S4.**
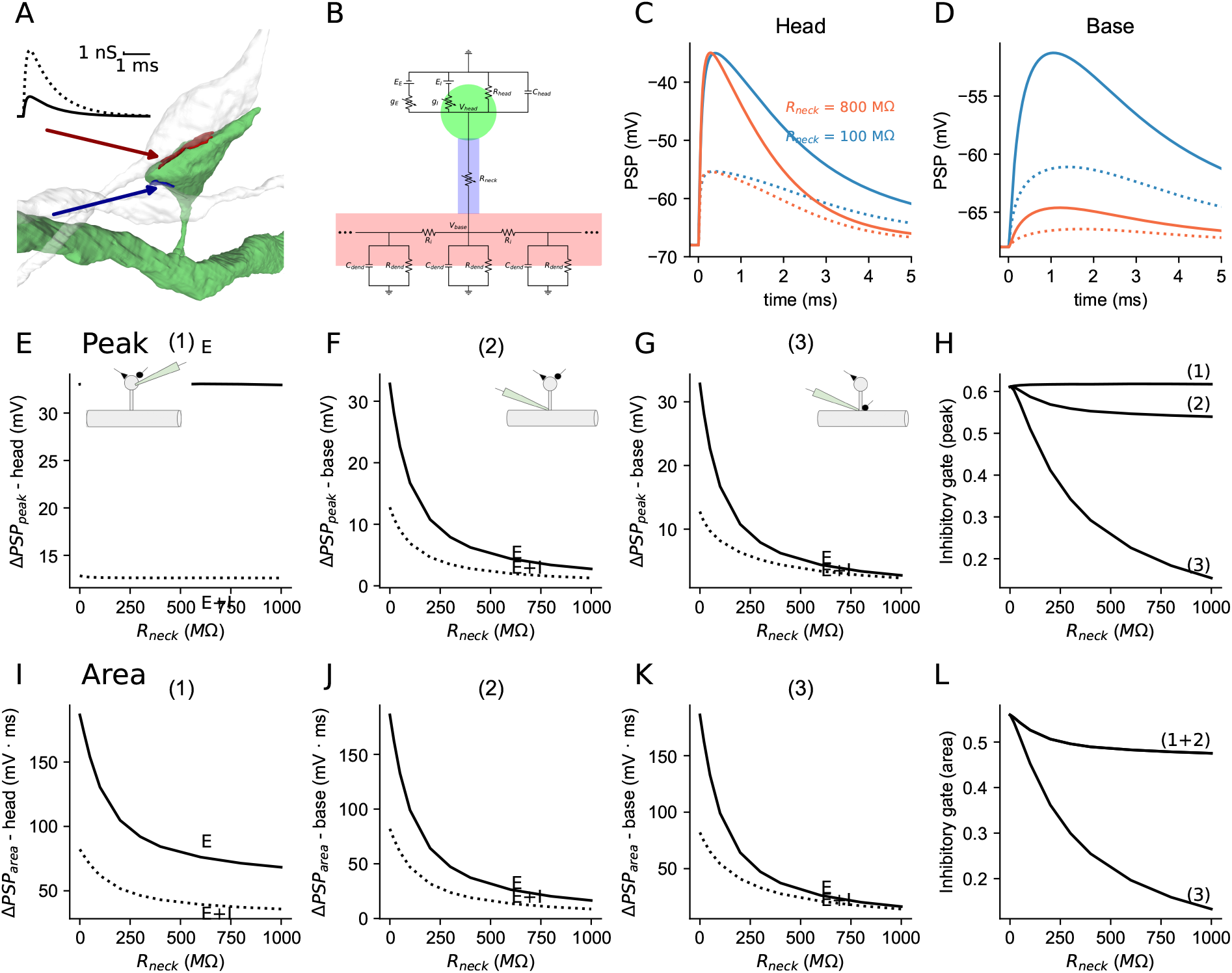
Potent inhibitory gating in dually innervated dendritic spines (DiSs) with −35 mV EPSP peak and AMPA-like inhibition. **A**. An EM-reconstruction of DiS. The two synapses are shown in red and blue (and with respective arrows). The inset shows the amplitude and time-course of the excitatory (continuous line) and inhibitory (dotted line) conductances. **B**. Equivalent circuit for a DiS on a near-infinite passive dendritic cable. **C**. PSPs at the spine head with (dotted line) and without inhibition (continuous line) for *R*_*neck*_ = 100*M* Ω (blue) and 800*M* Ω (orange). **D**. As in C, but the resultant PSPs at the spine base are shown. **E**. PSP peak at the spine head without (E) and with (E + I) inhibition as a function of *R*_*neck*_. **F**. As in E, but at the spine base. **G**. PSP peaks at the spine base when E is at the spine head and I at the spine base. **H**. Inhibitory gate (Equation 7) for the cases shown in E-G. **I-L**. Similar to E-H, but for area. Model parameters: AMPA: *τ*_*rise*_ = 0.1*ms, τ*_*decay*_ = 1*ms*. GABA: *τ*_*rise*_ = 0.1*ms, τ*_*decay*_ = 1*ms, E*_*GABA*_ = −68*mV* (equals the resting potential). *g*_*max*_ = 1*nS* for AMPA and GABA were automatically calculated such that the peak EPSP (AMPA only) will be equal to −35 mV peak.

**Figure S5.**
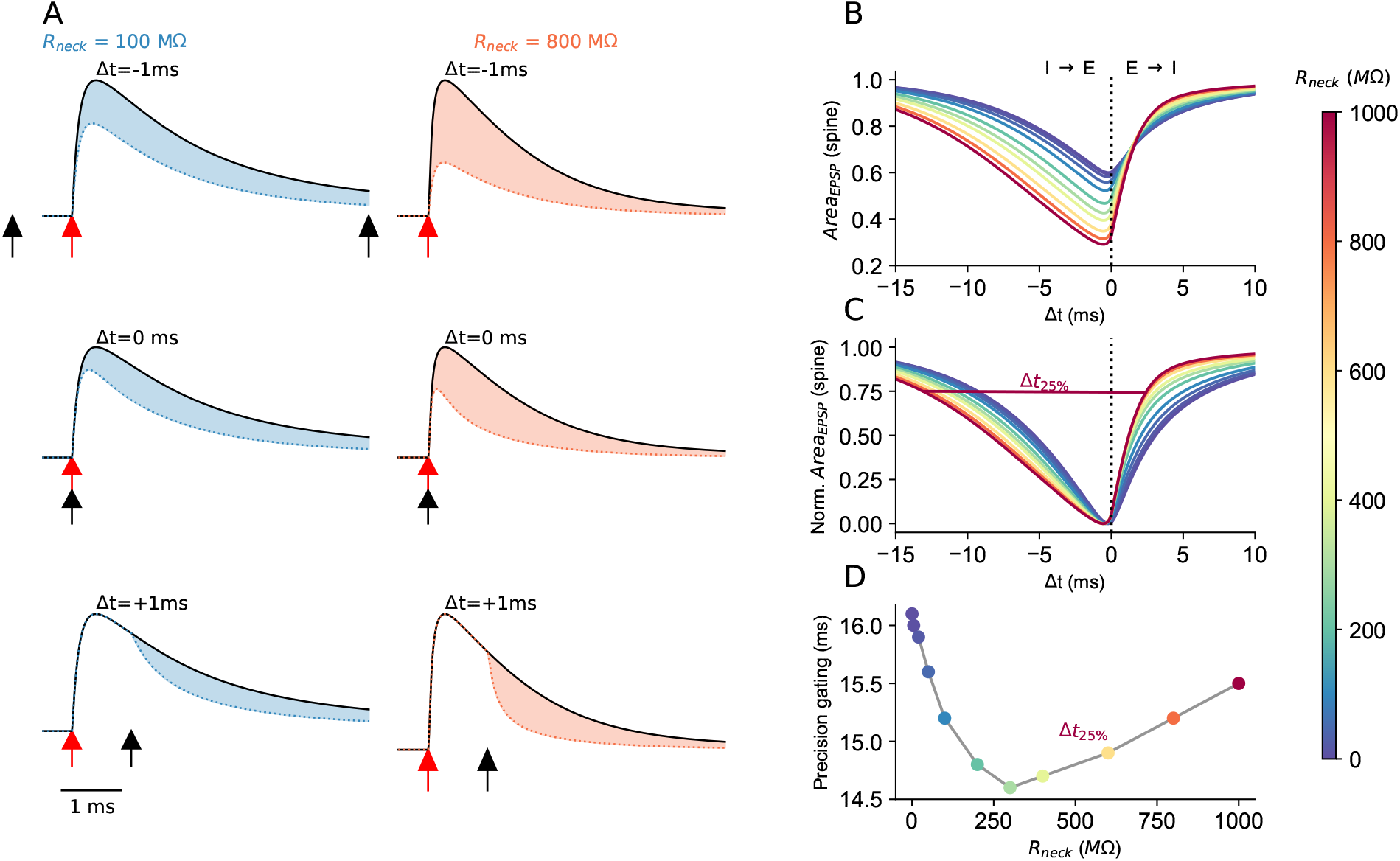
*R*_*neck*_-dependent temporal precision inhibitory gating on area of spinous EPSP in dually-innervated spines. **A**. Normalized PSPs at the spine head for *R*_*neck*_ = 100*M* Ω (blue) and 800*M* Ω (red). There are three cases of Δ*t* – the time difference between I (black arrow) and E (red arrow) activation. **B**. Normalized PSP area as a function of Δ*t* for various *R*_*neck*_ values (color code at right). **C**. As in B, but all curves are normalized to range between 0-1; the temporal precision gating of I over E (Δ*t*_25%_) is defined as the width of the curves at 25% reduction of the PSPs area. **D**. Δ*t*_25%_ as a function of *R*_*neck*_. The model is as in Figure S3.

**Figure S6.**
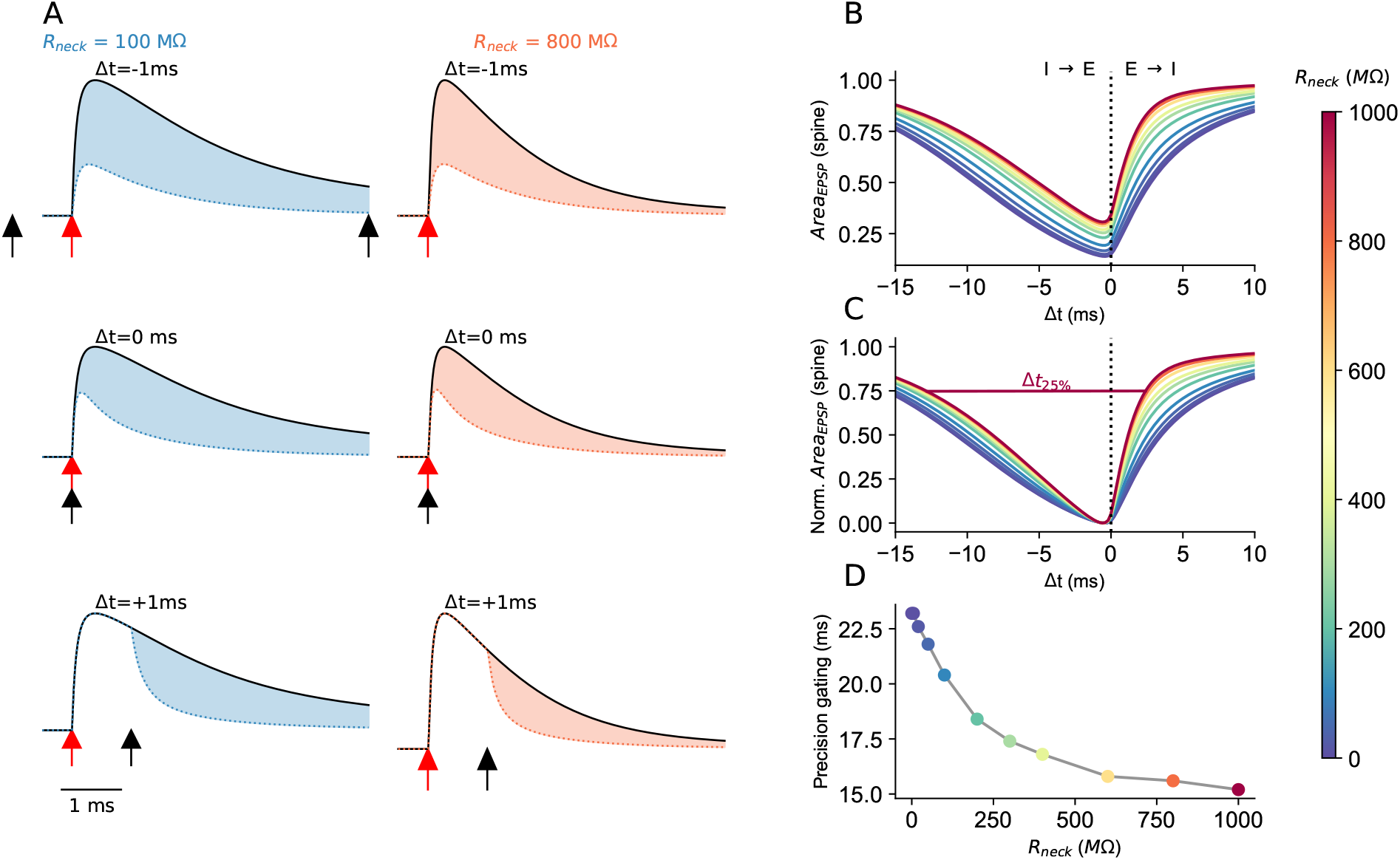
*R*_*neck*_-dependent temporal interactions between excitation and inhibition in dually-innervated spines. **A**. Normalized PSPs at the spine head for *R*_*neck*_ = 100*M* Ω (blue) and 800*M* Ω (red). There are three cases of Δ*t* – the time difference between I (black arrow) and E (red arrow) activation. **B**. Normalized PSP area as a function of Δ*t* for various *R*_*neck*_ values (color code at right). **C**. As in B, but all curves are normalized to range between 0-1; the temporal precision gating of I over E (Δ*t*_25%_) is defined as the width of the curves at 25% reduction of the PSPs area. **D**. Δ*t*_25%_ as a function of *R*_*neck*_. The model is as in Figure S3.

**Figure S7.**
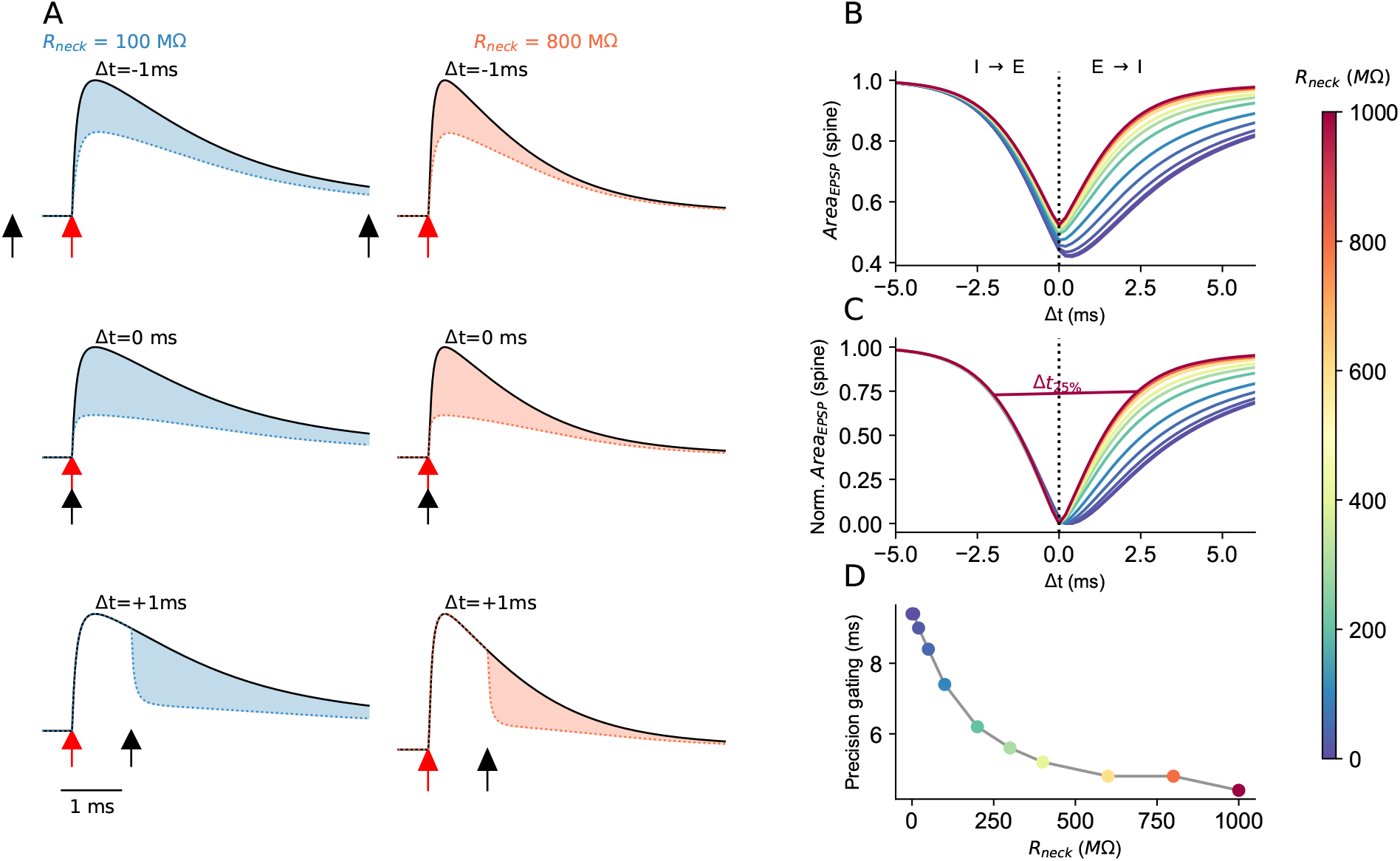
*R*_*neck*_-dependent temporal interactions between excitation and inhibition in dually-innervated spines and −35 mV EPSP peak and AMPA-like inhibition. **A**. Normalized PSPs at the spine head for *R*_*neck*_ = 100*M* Ω (blue) and 800*M* Ω (red). There are three cases of Δ*t* – the time difference between I (black arrow) and E (red arrow) activation. **B**. Normalized PSP area as a function of Δ*t* for various *R*_*neck*_ values (color code at right). **C**. As in B, but all curves are normalized to range between 0-1; the temporal precision gating of I over E (Δ*t*_25%_) is defined as the width of the curves at 25% reduction of the PSPs area. **D**. Δ*t*_25%_ as a function of *R*_*neck*_. The model is as in Figure S4.

